# Few-shot pattern detection by transient boosting of somato-dendritic coupling

**DOI:** 10.1101/2024.01.16.575776

**Authors:** Gaston Sivori, Tomoki Fukai

## Abstract

Neurons are thought to detect salient patterns amidst noise in continuous information streams, but their rapidity tends to be overlooked. Consequently, theoretical neuron models lack key mechanistic features that are suggested to underlie biological neuron rapid learning of input patterns. To unravel these features, we propose a class of models endowed with biologically-plausible predictive learning rules. In these models, an error signal propagates somatic spiking activity to dendrites, facilitating unsupervised learning of repeatedly coactivated presynaptic-neuron communities. Spike-triggered transient boosting of dendritic coupling bestows plausibility and improves the signal-to-noise ratio of learning dramatically. We demonstrate that our plasticity rule enables neurons to swiftly establish a behavioral timescale reward-place association in spatial navigation tasks and showcase how cell assemblies pre-configured in recurrent networks learn multiple patterns within a few repetitions robustly. Our results shed light on the self-supervising function of backpropagating action potentials for pattern learning and its acceleration by pre-existing cell assemblies.

---

Evidence indicates that cortical neurons can detect co-activation patterns^1,2^ and temporal sequences of presynaptic neurons^3^ in a stream of synaptic input. Since spike patterns may contain task-relevant information such as sensory stimuli^4^ and behavioral contexts^5^, computational models have been proposed to understand the mechanism of pattern detection in neurons^6–12^ and the roles of such patterns in brain computing^9,13,14^. However, despite extensive experimental and computational efforts^15–22^, it remains unclear how neurons rapidly become selective to specific patterns of synaptic inputs contaminated by biological noise. This may require extensive monitoring of all potential inputs to dendritic trees of individual neurons and their cooperative computing at the network level^10^.

Two different computational approaches can be made to pattern detection in neurons. In one approach, synaptic learning rules are derived from experimental evidence such as spike-timingdependent plasticity (STDP)^4,23–27^. For instance, using experimentally-tuned parameters for STDP in an unsupervised neuron model can show how these models become close to optimal in detecting coincidences^9,13,28^. Though such an approach can be informative, obtained results should be interpreted with care since the details of learning rules on which a particular model is based may depend on experimental conditions (culture compositions, single spikes or bursts, dendritic recording sites, etc.). An alternative approach is more mathematical, attempting to implement machine learning-based algorithms in biological neuron models. The latter tends to perform well on various tasks as the underlying learning rules are designed to optimize their performance in learning. Neuron models that use dendrites in implementing predictive learning rules are of particular interest as they can perform various pattern and sequence learning tasks^8,11^.

However, many of these biologically or mathematically well-based models require an unrealistic number of pattern presentations for learning. In addition, the mathematical models are computationally costly, rely on complex mathematical derivations, learn under non-biological assumptions, or fail to provide explanations of how they converge to their target output, among other weaknesses. Therefore, how biological neurons rapidly detect and integrate groups of synaptic inputs that repeatedly depolarize their dendritic tree remains elusive.

In this study, we present a novel synaptic plasticity rule that explains how principal cells can rapidly adapt their spiking response to temporal patterns of input in an unsupervised manner under biological assumptions and constraints. Our proposed biological rule builds on the machine-learning-based rule derived from the hypothesis that the dendritic synaptic activity learns to predict the somatic spike response^8,11,12,29^, broadening the biological plausibility of such prospective neural computing rules. We propose a Hebbian-like synaptic plasticity rule and test it in a set of two-compartmental neuron models that incorporate several biophysical mechanisms suggested to facilitate learning. We explore the learning paradigm by exposing the neuron models to a set of patterns repeating randomly and intermittently in afferent input. The task for our cell models is to pick up any of these statistically salient patterns hidden amidst the stream of synaptic input and produce robust spiking activity when it detects said pattern.

Furthermore, we demonstrate the performance of the proposed plasticity rule in two biologically realistic examples. In the first example, we test the ability of our learning rule to facilitate learning place fields in a realistic scenario in which synaptic inputs are initially non-structured and therefore the model neuron has no spatial receptive fields. During navigation, hippocampal neurons are known to rapidly develop selective responses to arbitrary spatial locations^30^. We examine the crucial role of N-methyl-D-aspartate receptor (NMDAR) temporal dynamics in promoting place field learning by deliberately blocking (i.e. silencing) and compensating for NMDAR-mediated Ca^2+^ influx when simulated mice run on a treadmill while receiving place and reward cue information. Our results shed light on the boosting effects of NMDARs on behavioral timescale learning through calcium-driven synaptic plasticity.

In the second example, we examine the ability of our plasticity rule to detect and learn temporal communities hidden in noisy input to a recurrent neural network. Evidence from sensory^31^, motor^32^, and memory processing ^33,34^ suggests the active role of pre-existing assemblies of cortical neurons in learning novel experiences. We embed tightly connected neuron assemblies into a recurrent network model. These cell assemblies give a reservoir of activity patterns available for encoding novel experiences and dramatically speed up the detection and learning of temporal communities in afferent inputs. Our results suggest that the predictive learning rule and pre-existing cell assemblies provide the neural substrate for one-shot learning of episodes.

## Results

Previous neuron models based on machine-learning algorithms suggested that learning in single cortical neurons is highly sensitive to statistically salient input patterns ^11,12,29^. Here, a neuron model implementing unsupervised minimization between somatic and dendritic activities is particularly interesting as it demonstrated self-supervised chunking of temporally structured spike inputs to perform various cognitive tasks^8^. Below, we implement biologically plausible mechanisms underlying such machine learning-based learning in a two-compartment neuron model and use it to clarify the conditions enabling the rapid detection of activity patterns (within several times of presentation), which was not the case in the machine-learning-based model.

### Spike-trace-based model for unsupervised predictive learning

Supervised^11^ and unsupervised^8^ predictive learning rules implemented in single neurons hypothesize that the dendritic activity driven by synaptic inputs attempts to predict the somatic response in a statistical sense. To this end, synaptic weights *w* on the dendrites undergo plastic changes to minimize errors (or information loss) between somatic and dendritic compartment activities. We approximately implement unsupervised predictive learning in a two-compartment model subject to a Hebbian-like learning rule (Fig. 1A), which modifies the weight of synapse *i* by using a “self-supervising” signal *PI*(*i, t*) given to the dendrite. The somatic compartment is a leaky integrate-and-fire unit, that is, whenever the somatic compartment membrane potential reaches a threshold, a spike is artificially produced and the neuron resets and enters a refractory state. We describe the core of our biological learning rule below. The self-supervising signal comprising an intrinsic error signal *e*(*t*) that entirely depends on the postsynaptic somatic response, postsynaptic potentials (PSPs) evoked by presynaptic spikes (Fig. 1B), and a weight scaling function *ζ* (|*w*_*i*_|) (Fig. 1C). The following equations 1 to 3 and Eqs. 14-17 (Methods) describe the proposed learning rule:

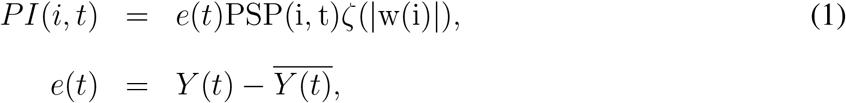

where *Y* (*t*) represents a graded spike count (decaying with a time constant *τ*_*Y*_) of somatic spike train *S*(*t*) = Σ _spikes_ *δ*(*t − t*_spikes_) and 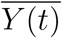 is the low-pass filtered spike trace obtained by solving Eqs. 2 and 3:

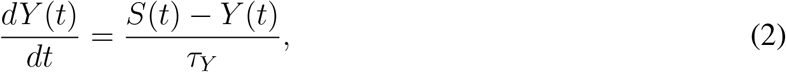

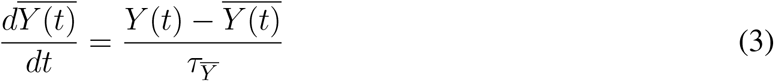

These spike traces are only utilized for the plasticity rule and their instantaneous values do not affect the membrane potential dynamics. The synaptic scaling term *ζ*(|*w*(*i*)|) comes from an Alpha function with a unitary range (Eq. 15 in Methods) and is determined to generate a synaptic strength distribution that matches the probability density of typical spine head volumes. Physiological spine growth is highly volatile and occurs relatively fast^35^, and spine volumes are a great proxy for synaptic strength distribution ^36^. Finally, before inducing long-term changes in synaptic weight *i*, we low-pass filter *PI*(*i, t*) with time decay constant *τ*_Δ_ and update the synaptic values at a learning rate *η* (Eq. 16 in Methods). Tables 1 and 2 summarize the values of the parameters for synapse and neuron models.

**Figure 1:**
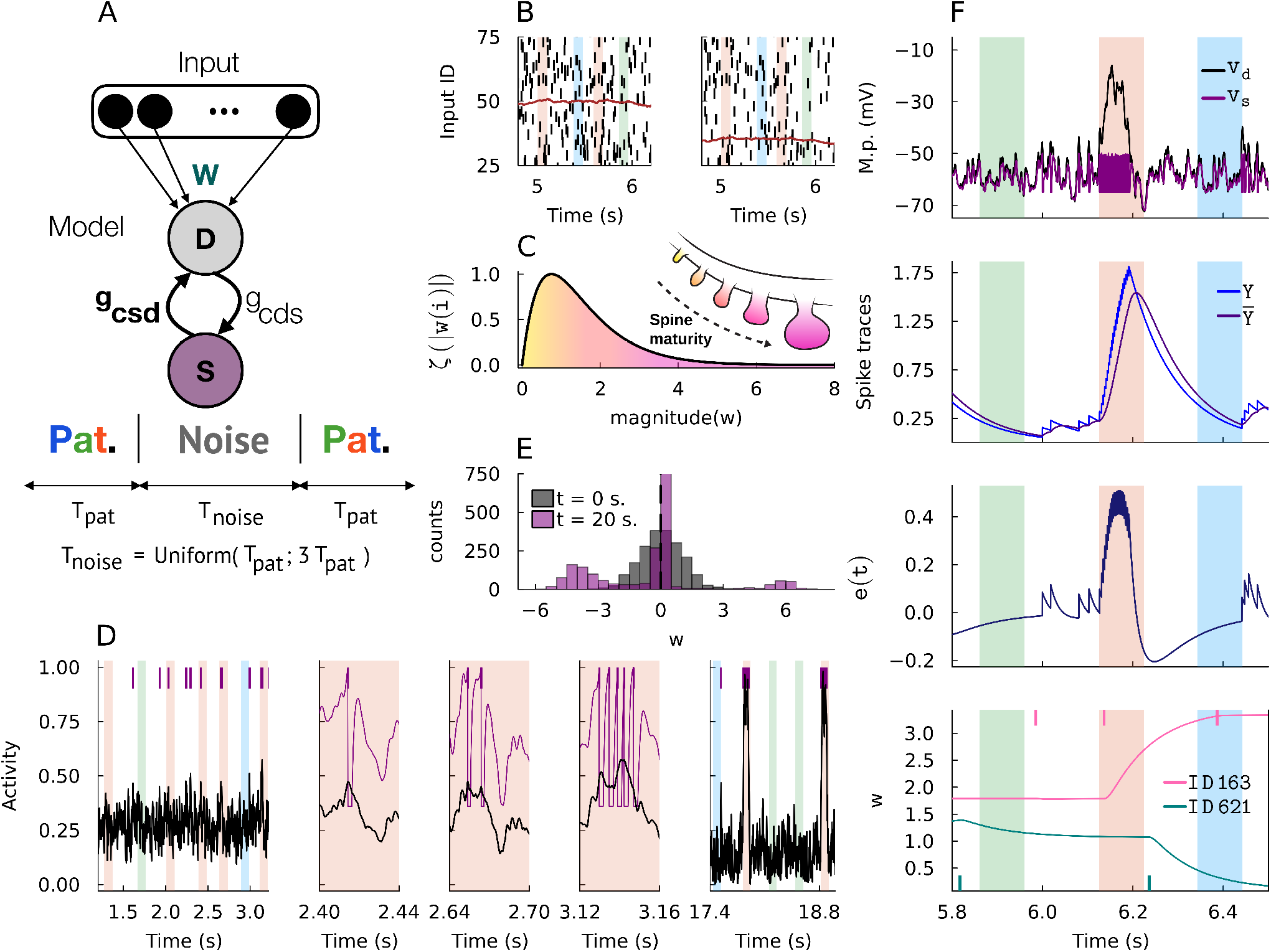
Spike-trace-based synaptic learning rule. (**A**) The two-compartment neuron model and input scheme. S: somatic compartment. D: dendritic compartment. Input: Poisson spiking units. w: synaptic strengths vector. *g*_*cds*_: dendro-somatic coupling conductance. *g*_*csd*_: somato-dendritic coupling conductance. Patterned spike inputs of length *T*_*pat*_ occur randomly in temporal input spike trains and are separated by noise spike patterns of length *T*_*noise*_ sampled from a uniform distribution ranging from *T*_*pat*_ to 3 *T*_*pat*_. (**B**) Examples of sent (left) and received (right) spike trains are shown. Due to transmission failure, these spikes trains have different mean spike counts (brown colored traces, same valued y-axis). (**C**) A *ζ*-scaling function was fitted phenomenologically based on spine maturity distributions. (**D**) Normalized dendritic activity (black) and somatic activity (purple spikes and traces) at different moments during a trial simulation. Note that somatic activity resets when it reaches threshold. (**E**) Synaptic weight distributions at start (t=0) and end (t=20) of a trial simulation.(**F**) Time courses of various dynamical variables of the spike-trace-based plasticity rule are shown. From top to bottom: the somatic and dendritic membrane potentials, spike trace *Y* (*t*) and its averaged (low-pass filtered) trace 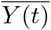, error signal *e*(*t*), and the weights *w* of two example synapses. Vertical colored bars indicate the periods of patterned inputs. Input spikes occurring during positive and negative values of *e*(*t*) are potentiated or depressed, respectively.

**Table 1:** Cell model parameter symbols, values, and descriptions.

| Symbol | Spike-based | Calcium-based | Ca<sup>2+</sup> and NMDA model | Description |
| --- | --- | --- | --- | --- |
| $g_{cds}$ | 108.0 | 108.0 | 108.0 | Dendro-somatic conductance in $nS$ |
| $g_{csd}$ | 8.0 | Equation 27 | Equation 27 | Somato-dendritic conductance in $nS$ |
| $g_{csdr}$ | $NA$ | 8.0 | 8.0 | Somato-dendritic resting conductance in $nS$ |
| $g_{ls}$ | 12.0 | 12.0 | 12.0 | Somatic leakage conductance in $nS$ |
| $g_{lsr}$ | $NA$ | 150.0 | 150.0 | Somatic leakage conductance during refractory period in $nS$ |
| $g_{ld}$ | 10.0 | 10.0 | 10.0 | Dendritic leakage conductance in $nS$ |
| $\overline{g_e}$ | 0.5 | 0.5 | 0.5 | Excitatory (AMPA) peak conductance in $nS$ |
| $\overline{g_i}$ | 0.55 | 0.55 | 0.55 | Inhibitory (GABA) peak conductance in $nS$ |
| $\overline{g_{Ca}}$ | $NA$ | 0.01 | 0.01 | HVA-Ca<sup>2+</sup> peak conductance in $nS$ |
| $\overline{g_{NMDA}}$ | $NA$ | $NA$ | 0.0005 | NMDA receptor peak conductance in $nS$ |
| $C_s$ | 180.0 | 180.0 | 180.0 | Somatic capacitance in $pF$ |
| $C_d$ | 60.0 | 60.0 | 60.0 | Dendritic capacitance in $pF$ |
| $V_{th}$ | -50.0 | -50.0 | -50.0 | Membrane threshold potential in $mV$ |
| $V_{peak}$ | $NA$ | 35.0 | 35.0 | Membrane peak potential in $mV$ |
| $V_{re}$ | -60.0 | $NA$ | $NA$ | Membrane reset potential in $mV$ |
| $V_{ls}$ | -70.0 | -70.0 | -70.0 | Somatic membrane resting potential in $mV$ |
| $V_{ld}$ | -70.0 | -70.0 | -70.0 | Dendritic membrane resting potential in $mV$ |
| $V_{er}$ | 0.0 | 0.0 | 0.0 | Excitatory reversal potential in $mV$ |
| $V_{ir}$ | -75.0 | -75.0 | -75.0 | Inhibitory reversal potential in $mV$ |
| $V_{Ca}$ | $NA$ | 120.0 | 120.0 | Ca<sup>2+</sup> reversal potential in $mV$ |
| $V_{NMDA}$ | $NA$ | $NA$ | 0.0 | NMDA receptor reversal potential in $mV$ |
| $\tau_{e, rise}$ | 0.5 | 0.5 | 0.5 | Time rise constant for excitatory synapses in $ms$ |
| $\tau_{e, decay}$ | 3.0 | 3.0 | 3.0 | Time decay constant for excitatory synapses in $ms$ |
| $\tau_{i, rise}$ | 1.0 | 1.0 | 1.0 | Time rise constant for inhibitory synapses in $ms$ |
| $\tau_{i, decay}$ | 8.0 | 8.0 | 8.0 | Time decay constant for inhibitory synapses in $ms$ |
| $\tau_{nmda, rise}$ | $NA$ | $NA$ | 3.3 | Time rise constant for NMDA synapses in $ms$ |
| $\tau_{nmda, decay}$ | $NA$ | $NA$ | 102.38 | Time decay constant for NMDA synapses in $ms$ |
| $\tau_{csd, rise}$ | $NA$ | 0.2 | 0.2 | Time rise constant for $g_{csd}$ boosting in $ms$ |
| $\tau_{csd, decay}$ | $NA$ | 1.5 | 1.5 | Time decay constant for $g_{csd}$ boosting in $ms$ |
| $f$ | $NA$ | $NA$ | 0.05 | NMDA-to-$I_{Ca}$ current factor. |

**Table 2:**
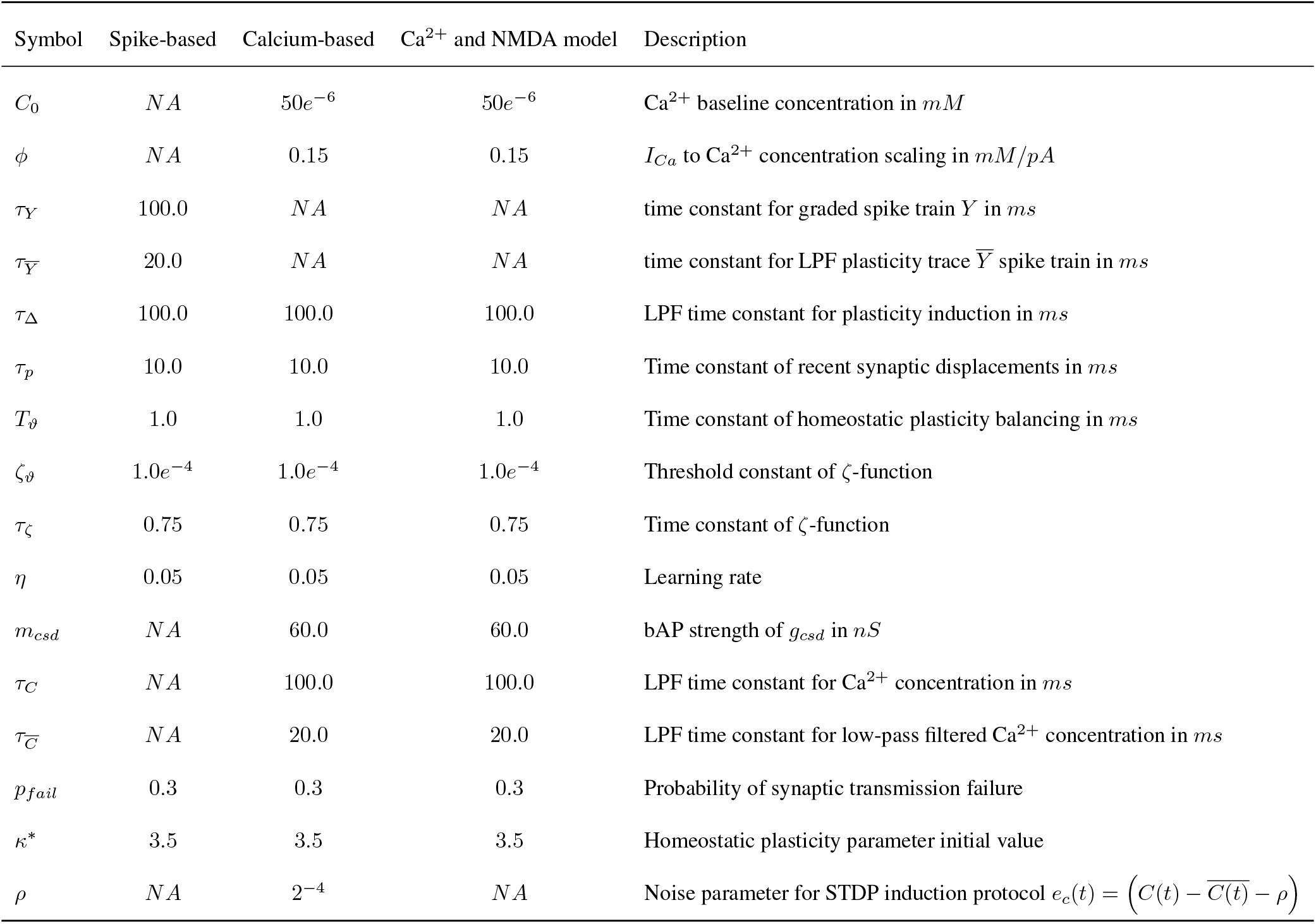
Synaptic plasticity rules’ parameter symbols, values, and descriptions.

| Symbol | Spike-based | Calcium-based | Ca<sup>2+</sup> and NMDA model | Description |
| --- | --- | --- | --- | --- |
| $C_0$ | *NA* | $50e^{-6}$ | $50e^{-6}$ | Ca<sup>2+</sup> baseline concentration in *mM* |
| $\phi$ | *NA* | 0.15 | 0.15 | $I_{Ca}$ to Ca<sup>2+</sup> concentration scaling in *mM/pA* |
| $\tau_Y$ | 100.0 | *NA* | *NA* | time constant for graded spike train *Y* in *ms* |
| $\tau_{\bar{Y}}$ | 20.0 | *NA* | *NA* | time constant for LPF plasticity trace $\bar{Y}$ spike train in *ms* |
| $\tau_{\Delta}$ | 100.0 | 100.0 | 100.0 | LPF time constant for plasticity induction in *ms* |
| $\tau_p$ | 10.0 | 10.0 | 10.0 | Time constant of recent synaptic displacements in *ms* |
| $T_{\vartheta}$ | 1.0 | 1.0 | 1.0 | Time constant of homeostatic plasticity balancing in *ms* |
| $\zeta_{\vartheta}$ | $1.0e^{-4}$ | $1.0e^{-4}$ | $1.0e^{-4}$ | Threshold constant of ζ-function |
| $\tau_{\zeta}$ | 0.75 | 0.75 | 0.75 | Time constant of ζ-function |
| $\eta$ | 0.05 | 0.05 | 0.05 | Learning rate |
| $m_{csd}$ | *NA* | 60.0 | 60.0 | bAP strength of $g_{csd}$ in *nS* |
| $\tau_C$ | *NA* | 100.0 | 100.0 | LPF time constant for Ca<sup>2+</sup> concentration in *ms* |
| $\tau_{\bar{C}}$ | *NA* | 20.0 | 20.0 | LPF time constant for low-pass filtered Ca<sup>2+</sup> concentration in *ms* |
| $p_{fail}$ | 0.3 | 0.3 | 0.3 | Probability of synaptic transmission failure |
| $\kappa^*$ | 3.5 | 3.5 | 3.5 | Homeostatic plasticity parameter initial value |
| $\rho$ | *NA* | $2^{-4}$ | *NA* | Noise parameter for STDP induction protocol $e_c(t) = (C(t) - \overline{C(t)} - \rho)$ |

Unless otherwise stated, throughout this study, we test the performance of our model in a temporal pattern detection task depicted in Fig. 1A. An input layer consists of 2000 neurons (approximately 1000 excitatory and 1000 inhibitory units), all of which generate inhomogenous Poisson spike trains. The afferent inputs have a duration of 20 seconds and involve repeated spike patterns of multiple assemblies of neurons (color-shaded domains in Fig. 1B). Each assembly consists of about 230 excitatory and inhibitory neurons and is mutually non-overlapping. In this study, we use three repeated patterns, although increasing the number of patterns does not change the essential results. Consecutive repeated patterns are separated by non-repeated random spike trains (non-shaded domains in Fig. 1B) of which duration is sampled uniformly, as shown in Fig. 1A. The repeated patterns have an identical duration and occur with equal probabilities.

Our learning rule rapidly increases the detection capacity of repeated structured input, as demonstrated in Fig. 1D showing changes in the output spike count of the model neuron before, during, and after an example trial. The model neuron eventually learned a selective response to one of the repeated patterns. There was no explicit target pattern for the neuron model, but the nature of the learning rule allowed it to pick up any of the statistically salient patterns hidden in the input stream. Surprisingly, only several times of exposure to the pattern sufficed for this learning. We regarded that convergence was reached when the model was capable of responding robustly to said pattern. When pattern templates were created randomly, the neuron model developed either an early onset response or late onset response within the pattern time and, in both cases, satisfied our pattern detection criteria.

The weight distributions of excitatory and inhibitory synapses obtained at the end of the simulation displayed bimodality (Fig. 1E). The strong selectivity of the learned neuronal response is hinted from strong excitatory synapses mediating the detected input pattern and strong inhibitory synapses mediating non-detected input patterns. When combined, these distributions yield a longtailed distribution with dense weak links and sparse strong links. Cortical synapses are known to obey long-tailed strength distributions^37^ and their computational implications have been studied extensively in memory encoding and retrieval^38^. In our model, the growth of synapses is controlled by the scaling function that is multiplicative with the Hebbian term (see Fig. 1C and Eq. 15 in Methods). Pattern selectivity is further hinted in synaptic distributions of pattern synapses across different time snapshots during training (Supplementary Fig. 1).

### Relationship to machine-learning-based learning rules

In the original machine-learning approach^11^, synaptic plasticity is governed by an instantaneous prediction error, 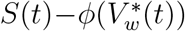, which is given by the somatic spike train *S*(*t*) (driven externally by a teaching signal) and dendritic prediction of the actual somatic firing, 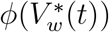, calculated from the rescaled dendritic membrane potential 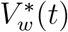 through a non-linear filter *ϕ*. Similarly, we can derive an unsupervised learning model from the supervised one by removing the teaching signal and introducing an additional mechanism to regulate the cell’s intrinsic excitability^8^. By construction, these models are successful in learning non-random, hence predictable, statistical features (e.g., repeated spike patterns) hidden in synaptic input. However, they are built on mathematical assumptions of which biological reality is unclear, and several questions remain to be answered. Can single neurons calculate the probability distributions of somatic and dendritic activities and the distance between them, which underlie the predictive learning rule? What biological mechanisms are required for this? More specifically, the previous models assume that the dendrite has a readily available signal that predicts somatic output. Such a signal is particularly crucial for unsupervised learning as it solely relies on it. Finally, are thus-constructed biological neuron models equally (or even more) efficient in detecting and learning non-random input features?

The learning rule proposed in this study aims to address these questions. Moreover, while sharing certain similarities, the present plasticity rule can solve the difficulty encountered by the machine-learning-based unsupervised learning rule. Unlike the machine-learning-based rule that relied on a memory trace of presynaptic inputs, our error signal depends on a memory trace 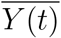 of recent postsynaptic spikes. The dependency on the history of synaptic inputs or that of postsynaptic firing forces the neuron’s current spiking activity explicitly or implicitly to obey the probabilistic structure of learned salient synaptic inputs. Thus, these memory traces represent the predictive characteristic of the learning rules. However, the machine-learning-based unsupervised rule^8^ struggled with the trivial solution of all weights converging to zero (*w →* 0), because this gives the easiest case where no spiking output is predicted for no synaptic input. In contrast, our biological rule can avoid this difficulty by producing sustained fluctuations around a baseline output activity. At any given time when *e*(*t*) is positive, all inputs that contributed to spiking are potentiated by a sampled fraction of the error signal. When *e*(*t*) becomes negative (typically when spiking has recently ceased), any input after it will be depressed. Thus, the error signal serves as a proxy for recent variations of postsynaptic activity. The amounts of synaptic plastic changes are eventually balanced between long-term potentiation (LTP) and long-term depression (LTD) when the areas under the curve (positive and negative) *e*(*t*) become equivalent.

### Facilitation of temporal community detection by transient boosting of somato-dendritic coupling

The results shown in Fig. 1 demonstrated that the proposed learning rule enables the two-compartmental neuron to detect temporal activity patterns rapidly. To accelerate the pattern detection, this learning rule uses a plasticity signal that is updated instantaneously for every somatic spike and becomes available at all active synapses, as in STDP, which gives a convenient mechanism of pattern detection^9,13,28,39^. However, such a signal is not readily available in biological cells. Therefore, we now construct a slightly more plausible model that takes into account the mechanistic function of backpropagating action potentials (bAPs)^40,41^.

In the following model, we introduce high-voltage activated (HVA) Ca^2+^ channel dynamics^42^ in the dendritic compartment (see Methods for channel equations) and an intracellular Ca^2+^ concentration trace via integration of calcium current *I*_*Ca*_ (Eq. 4). It is known that postsynaptic Ca^2+^ concentration dynamics are a proxy of somatic activity, especially in the basal and in tuft dendrites^43^, and have a pivotal role in synaptic plasticity^44^:

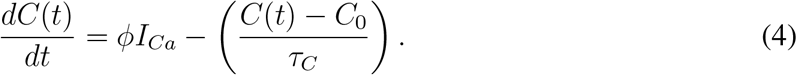

The parameter *ϕ* scales *I*_*Ca*_ into Ca^2+^ concentration, *C*_0_ is the Ca^2+^ concentration at rest, and *τ*_*C*_ is the time constant of Ca^2+^. We swap this *C*(*t*) trace for the spike train trace *Y* (*t*) in the synaptic plasticity rule. Similarly as in the previous rule, calculating an averaged trace 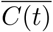 by low-pass filtering Ca^2+^ concentration, we obtain a modified error signal for our plasticity rule shown in equation (5):

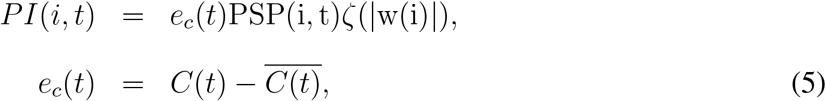

The rest of the equations (Eqs. 15-17 in Methods) in the plasticity rule remain unchanged.

Plugging in HVA-Ca^2+^ channel dynamics into the dendritic compartment is relatively straightforward and its effect on dendritic depolarization is minimal (see Table 1 for parameter values). With this change, synaptic plasticity updates can now occur, though small, even in the absence of postsynaptic spiking. However, the internal error *e*_*c*_(*t*) is not directly linked to somatic spiking activity and fails to reach a target pattern. We showcase an example failure trial where, compared to the spike-based model (Eq. 1), the calcium-based plasticity alone is unable to converge (Fig. 2A). Thus, the calcium-based plasticity mechanism alone is less efficient than the naive mechanism having direct access to the somatic spiking.

**Figure 2:**
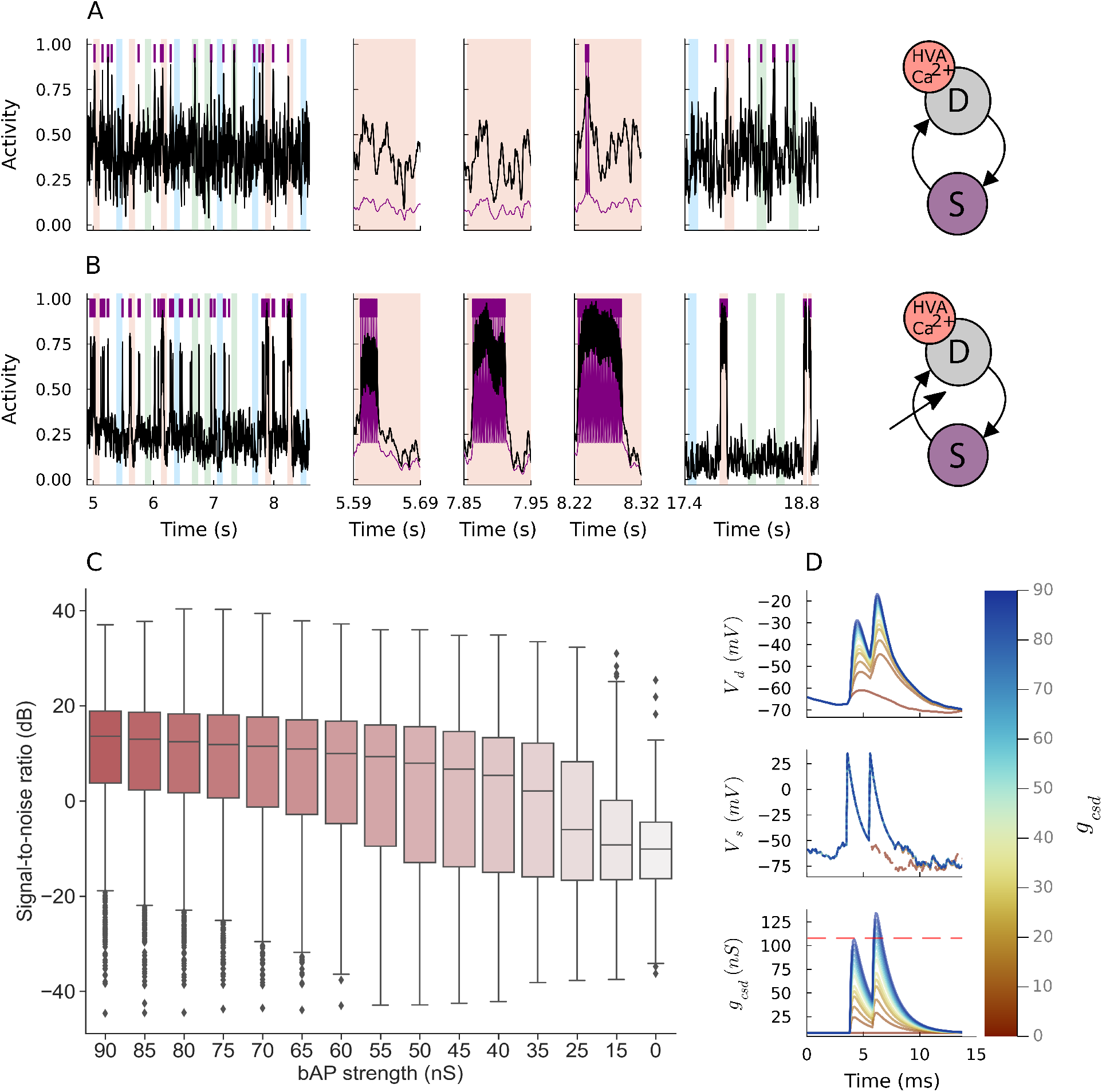
Calcium-based plasticity rule with spike-triggered transient somato-dendritic coupling boosting. (**A**) Normalized dendritic activity (black) and somatic activity (purple spikes and traces) at different moments during a trial simulation are shown for the calcium-based neuron model without a transient boosting of the somato-dendritic coupling. (**B**) Same as above but the neuron model includes transient boosting of somato-dendritic coupling during spike generation (the bAP strength of 60 nS). A cross arrow schematically illustrates the transient boosting of the somato-dendrite coupling. (**C**) Box plots containing five-number summary and outliers of signal-to-noise results (500 trials for each case; failure trials discarded) are shown for the Ca-based neuron model with the transient coupling boosting at different bAP strengths. (**D**) Effect of different bAP strength on actual AP backpropagation is assessed. Colors indicate each bAP strength and dashed red line is the dendrite-to-soma coupling. At bAP strength of 15 nS, the dendritic depolarization triggers a second somatic spike.

We introduce a transient boosting in somato-dendritic coupling to rescue the Ca^2+^-based model and recuperate its learning capacity (Fig. 2B). During a backpropagating event in biological cells, a wave of depolarizing current travels across the dendritic tree. We mimic this biophysical phenomenon by transiently increasing the coupling conductance from somatic to dendritic compartments (i.e., *g*_*csd*_ in Methods Eqs. 26-29). This key behavior only occurs during somatic spiking and is abrupt enough to avoid noisy somatic currents in the dendrite from sub-threshold somatic fluctuations. Namely, a large constant somatodendritic coupling would not work as it may also amplify somatic spiking caused by noisy input. As shown in Fig. 2B, the transient boosting in *g*_*csd*_ dramatically improves both speed and signal-to-noise ratio (SNR) of pattern detection by the Ca^2+^-based model. The result of the transient increase is an intrinsic positive feedback formed by the somato-dendritic coupling dynamics and dendritic Ca^2+^ dynamics. With somatic spiking, a transiently boosted somato-dendritic coupling further depolarizes the dendritic membrane potential. This causes an influx of calcium current through the HVA-Ca^2+^ channel, serving an increase of Ca^2+^ concentration that raises the magnitude of plasticity updates. We note that the upswing dendritic membrane potential makes calcium-based plasticity traces resemble *Y* (*t*) and 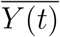.

As far as the somato-dendritic coupling generates sufficiently strong bAPs, our proposed rule with spike-based signals shows excellent learning performance. We performed 500 trial simulations and assessed the model’ performance by calculating the SNR at different bAP strength (measured by *m*_*csd*_ given in Methods Eqs. 28 and 29). We measured the signal as the number of spikes occurring during the tuned pattern presentation and noise as otherwise. The results presented in Fig. 2C indicate that the higher the bAP pulse, the stronger the performance of the model and its capacity for detection. However, the induced dendritic depolarization can grow out of proportion for extreme values of bAP pulse, increasing the number of outliers in the low SNR regime and, sometimes, even surpassing the opposite conductance of the somato-dendritic coupling (Fig. 2D). This implies that there exists an optimal range of the bAP pulse value for robust performance. If this value is too small, pattern detection becomes difficult, as demonstrated in Fig. 2A for an extreme case of vanishing somato-dendritic coupling. However, if the value is too large, even somatic spike responses to non-target patterns propagate to the dendrite and drive learning. Our model suggests that, in the biological scenario, a realistic bAP event is determined to solve this trade-off.

Excitatory and inhibitory conductances behave in a way that facilitates selective responses to a given pattern. In the example cases presented in Figs. 1 and 2 (bAP strength 60 nS), excitatory conductance remains high relative to inhibitory conductance during the presentation of pattern red (Supplementary Fig. 2A and 2C), whereas inhibitory conductance overwhelms the excitatory one during the presentation of the other patterns (Fig. 2B and 2D), resulting in strong red pattern selectivity. The transient somato-dendritic coupling increase forces the dendritic membrane potential upswings given somatic spiking, which introduces further somatic depolarization post refractoriness.

The choice of including the HVA-Ca^2+^ channel in our model is based on its activation threshold dynamics. There are, however, many other calcium channels that lead to further influx at different depolarization levels and may be useful to incorporate. However, our approach here only regards necessary mechanisms for producing rapid learning of hidden patterns amidst noise. It is an intriguing open question whether including other calcium channels can produce a similar effect to the transient somato-dendritic coupling increase on the pattern detection of the model.

### Crucial roles of NMDARs for rapid binding of patterns with low SNR

One crucial component that has a strong influence on the long-term potentiation and long-term depression of coincidently active synapses is the activation of NMDARs in dendritic spines^45^. An increased number of NMDA binding sites suggests a strengthened synapse, whereas a decrease number of binding sites is a sign of a weakened synapse^46^. The role of NMDARs in discriminating spatiotemporal activity patterns was also reported experimentally^47^. We investigate how the inclusion of NMDAR dynamics^48^ in the dendrite, which introduces sustained dendritic membrane potentials due to their slow timescale, affects the performance of our model. NMDARs also express permeability to calcium, and we model this by factoring out 5% of the total NMDA current to *I*_*ca*_ (see Methods equation 30).

In Fig. 3A we showcase an example trial for our Ca^2+^/NMDAR-based model. The model now includes HVA-Ca^2+^ channel equations, NMDAR equations, and the variant learning rule with calcium-based plasticity traces (Equation 5) and with the spike-triggered transient boosting of *g*_*csd*_. As expected, the neuron model rapidly converges to robust spiking to pattern red. For this case, we observe that the inclusion of NMDA receptor dynamics in the dendritic compartment can result in steeper rising and falling phases of the plasticity signal (Eq. 5), which now captures many more synapses during post-synaptic spiking. However, a question arises: how is the inclusion of NMDARs affecting the neuron’s pattern detection capacity?

**Figure 3:**
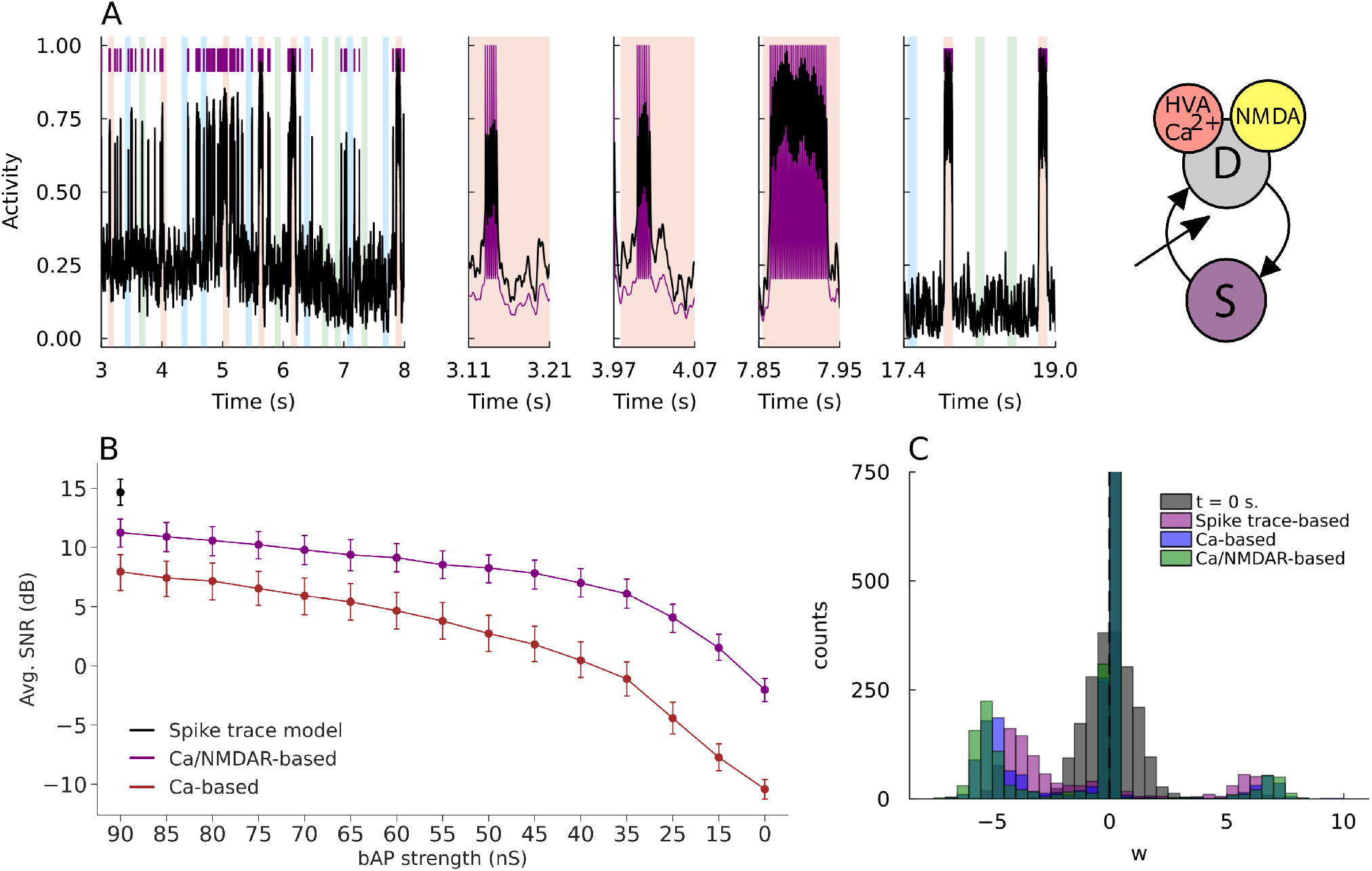
Enhanced detection capability of Ca^2+^-based plasticity rules with NMDA receptors. (**A**) Normalized dendritic (black) and somatic (purple) activity and spikes at different moments during trial simulation for calciumbased model including NMDAR dynamics. Panels from left to right depict early trial time, three red pattern presentations, and late trial time. Illustration for Ca-based neuron model including NMDA receptor dynamics and spiketriggered transient boosting of the somato-dendritic coupling. (**B**) Point plot of average SNRs of all models resulting from 500 simulation trials (no tuning trials discarded) at different bAP strengths. Error bars indicate confidence intervals (95%). Results of the spike trace-based model are plotted for comparison. Across all values of the bAP strength, the average SNR is significantly higher for the model including NMDAR dynamics. (**C**) Synaptic weight distributions at the start and end of trial simulations for all three models with the bAP strength of 60 [nS].

To answer this question, we calculated SNRs for each of the three different models, i.e., spike-trace-based, Ca^2+^-based, and Ca^2+^/NMDAR-based models, where the SNR is defined as in Fig. 2C and the latter two models employed the transient somato-dendritic coupling boosting. We calculated SNRs for a range of the bAP strength. For all models, the same learning rate (*η* in Methods Eq. 16) was utilized. We plot the average SNR in Figure 3B. The spike-trace-based plasticity rule yields the highest SNR among the three models. More importantly, the SNR of the calcium-based model with NMDAR dynamics (Ca^2+^/NMDAR-based) is nearly matched with the highest SNR. Thus, this model is the most capable at rapidly detecting patterns whilst remaining biologically plausible. As shown in Fig. 3C, in general, all models exhibit similar distributions of excitatory and inhibitory synaptic weights at the end of a given trial simulation where convergence was achieved. However, in the distribution of the Ca^2+^/NMDAR-based model, the peaks of strong excitatory and inhibitory synaptic weights are most clearly distinct from those of weak synapses, consistently with the excellent performance of this model.

### Robustness of pattern tuning against temporal structure corruption

In general, learning converges relatively fast within the present simulations, requiring a small number of pattern presentations. However, the learning efficiency of the model depends on various factors, including the number of coincident spikes within each pattern, the initial synaptic weight values, and noise intensity in input patterns. We explored how the model tunes to different patterns in the presence of spike timing jitters, varying the noise intensity. Input spike trains involved three different patterns (named blue, red, and green) with equal occurrence probabilities. Given enough simulation time, the model tuned to a specific pattern almost with equal probability (Fig. 4A). For many of these trials, a small number of presentations of the pattern were sufficient to reach convergence (notice graphs on shorter simulation times). In a small fraction of trials, the neuron model tuned to more than one input pattern, and the fraction of trials remained constant as the simulation time was prolonged (“Others” in Fig. 4A). This mixed tuning was presumably due to accidental overlaps among input patterns or unfavorable starting conditions.

**Figure 4:**
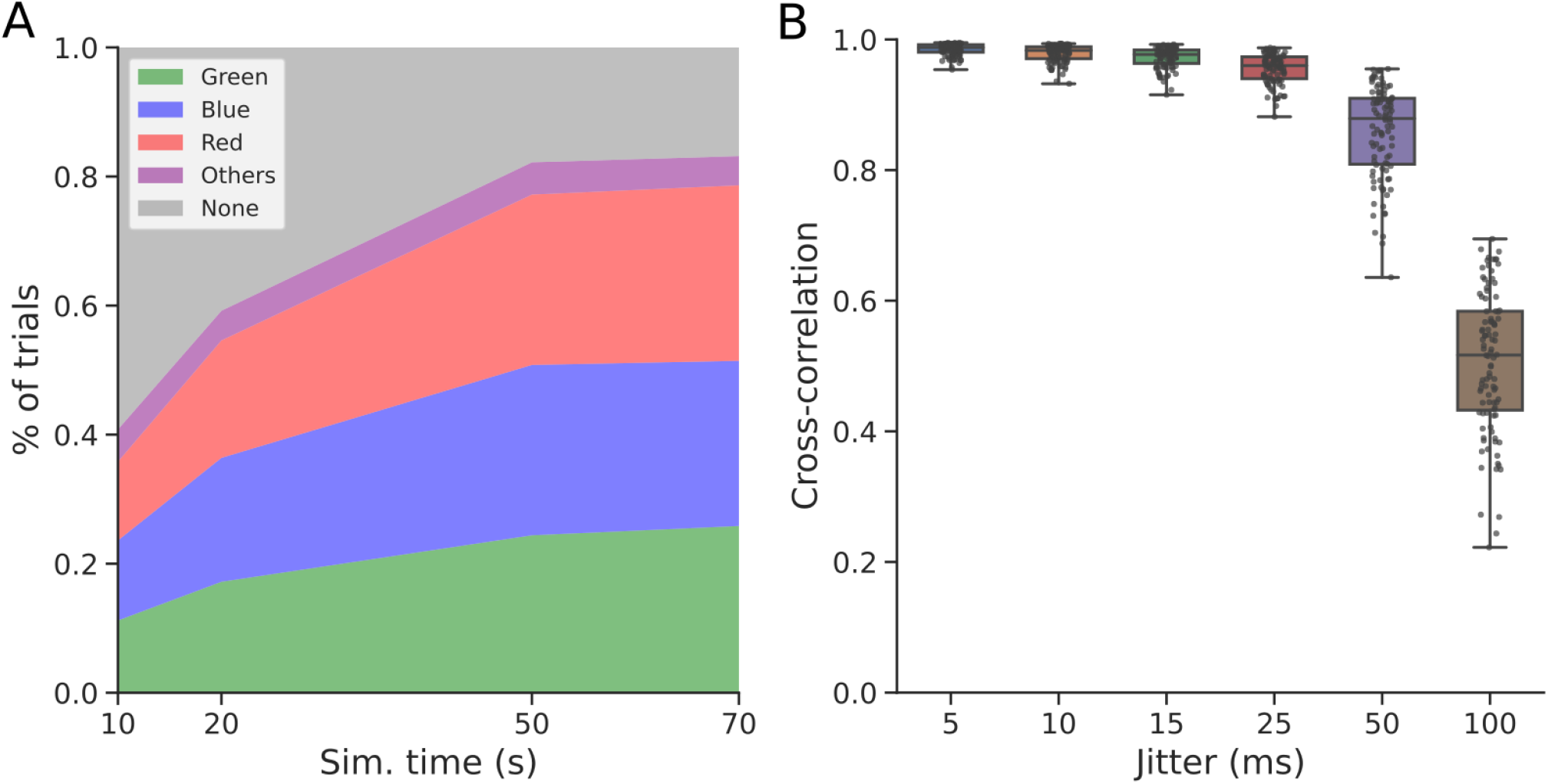
Convergence and robustness of pattern tuning against temporal structure corruption via uniform jittering. (**A**) The average percentages of trials in which the model tuned to particular patterns are plotted for simulation periods of 10, 20, 50, and 70 seconds. Five hundreds trials were performed in each case. “Others” label trials wherein more than one pattern was learned by the neuron. (**B**) Box plot overlayed with dot plot of cross-correlations between somatic spiking outputs given uniformly jittered input and the ground truth output are shown for different mean jitters. The data points on each box correspond to each 20-second simulation where model tuning for a given pattern converged. Box plots show five-number summary measures for each jitter case.

We highlight the importance of the temporal community structure within input patterns (i.e. temporally structured groups of coincident spikes). While the balance of excitation and inhibition (E/I) varies across each pattern presentation due to synaptic transmission failure (*p*_*fail*_), the temporal community belonging to a certain pattern is what governs the learning process. To study how robust the post-learning model performance is, we increased the mean jitter timing of input spikes (Fig. 4B). Namely, we selected the results of 100 previous 20-second simulation trials wherein pattern tuning was achieved, and evaluated the robustness of the model’s responses against the corruption of the original patterns. We tested the robustness by using a “ground truth” spiking output, which is a response of the post-learning neuron model to a “clean” input pattern without timing jitters. To prevent synaptic weights from adapting to noisy input patterns, we continued to freeze synaptic weights during the test and repeatedly applied noisy input patterns. We calculated cross-correlations between the ground truth output and outputs to the noisy patterns, revealing that pattern tuning is robustly maintained even for a 50-millisecond mean jitter but drops to a chance level for a 100-millisecond mean jitter, which coincides with pattern duration.

### Burst-induced Schaffer-collateral STDP replicated by the calcium-based model

STDP protocols have revealed important information about the plastic changes in incoming synapses on the postsynaptic neuron. It is widely accepted that protocols involving a single presynaptic spike and a single postsynaptic spike leads to LTP and LTD at excitatory cortical synapses^23,49,50^(see Shoueval et al.^26^ for review), depending on the relative times of these spikes. However, single spikes may not be sufficient for robust information transfer between in-vivo pyramidal neurons, hence STDP rules with bursts of spikes have also been studied. It was shown that CA3-to-CA1 excitatory synapses in the hippocampus exhibit Mexican-hat-shaped weight changes when a burst of 2 postsynaptic spikes were repeatedly paired with a presynaptic spike at a frequency of 5 Hz^50^. Namely, LTP and LTD were induced for shorter and longer relative times, respectively. Owing to postsynaptic spike bursts and different experimental preparations (see Discussion for the latter point), this protocol is considered to induce stronger bAPs in CA1 pyramidal cells. Below, we show that our calcium-based rule replicates this STDP rule without assistance from NMDARs. For comparison reasons, we followed the pairing protocols used in the experiment, with a total of 20-30 or 70-100 times of repetitions. We also adopted the convention used in that study, defining Δ*t* as the time interval between postsynaptic bursts and presynaptic spikes (Fig. 5B and C, inset).

**Figure 5:**
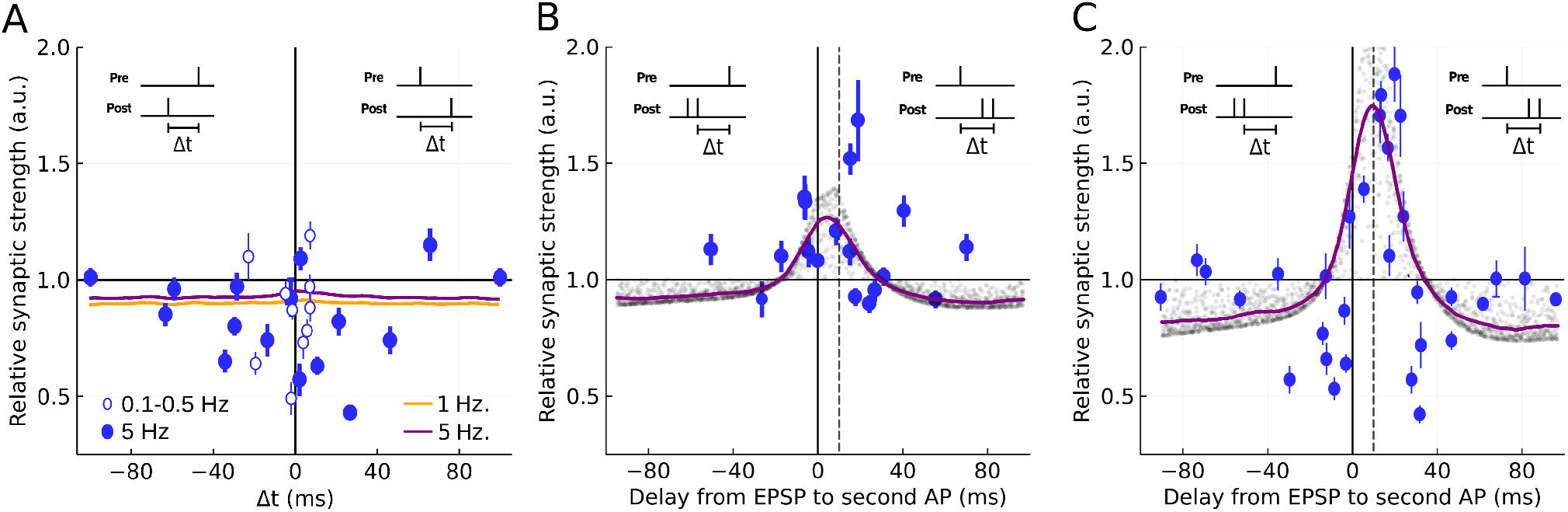
STDP rules obtained by our Ca^2+^-based plasticity rule. Each trial has many repetitions at different time delays between a presynaptic spike and a second somatic AP (inset), where positive intervals are defined as causal, and negative intervals as anti-causal. Simulation data points are presented with transparency to indicate time-window shapes. Colored solid lines are smoothed polynomial interpolations of simulation data points by linear least square fitting. Experimental data are shown with error bars indicating confidence intervals for 5 Hz (filled circles) and 0.10.5 Hz (open circles) pairing protocols. These data points are included with permission from the authors^50^. (**A**) The STDP rule of the model is shown for 70-100 pairings of singlet preand post-synaptic spikes at 5 Hz (purple) and 1 Hz (orange). Simulation data points are not included for visibility. (**B**) The model’s STDP rule is shown for 20-30 pairings at 5 Hz of presynaptic spikes and postsynaptic spike doublets. (**C**) The model’s STDP rule of the model is shown for 70-100 pairings at 5 Hz of presynaptic spikes and postsynaptic spike doublets.

In Figure 5, we show the STDP rules obtained from numerical simulations of the calcium-based model with experimental data points overlayed ^50^. To begin with, we examined the STDP rule resultant from 70-100 pairings of single presynaptic and postsynaptic spikes at frequencies of 1 Hz and 5 Hz. We introduced small noise in the model (parameter *ρ* in Table 2) to produce basal fluctuations in the error signal *e*(*t*), which result in a minimal LTD across all Δ*t* values (Fig. 5A). The experimental results suggest that the relative change in the synaptic strength becomes largest around small time differences (*−*40 ≲ Δ*t* ≲ 40 in ms). Running said protocol with our proposed rule, the average relative weight change displayed a slight increase around small time differences for both frequencies. However, since the increase was small and experimental data were also scattered, we cannot safely conclude that the results are consistent between experiments and numerical simulations.

In contrast, the simulation of the model neuron for bursts of postsynaptic spikes generates results more consistent with the experiment. When a single presynaptic spike was repetitively paired with a burst-like pair of postsynaptic spikes 20-30 times, pyramidal cells exhibited an LTP-only (or LTP-dominant) time window (Fig. 5B). The peak amplitude of LTP occurred near the origin of the time difference, where the spike singlet-doublet pairing protocol shifted the peak position slightly towards the positive (causal) side. Increasing the number of repetitions (70-100 pairings) enhanced the peak LTP amplitude, flanked by LTD sides that returned to baseline fluctuations towards longer time differences (Fig. 5C). Testing our proposed rule with this particular pairing protocol produces similar STDP time windows for both low and high numbers of repetitions. These results demonstrate that the proposed model describes a biologically plausible learning mechanism of cortical cells.

In vitro experiments have suggested that the LTD induced by causal pairings of presynaptic and postsynaptic spikes may be attributable to the presence of inhibition^24,25^. In the present simulations, the LTD sides that flank the LTP time window do not well fit the experimental data points. We suspect that the causes of these LTD time windows are related to biological mechanisms that have yet to be included in our model. These mechanisms seem to require a higher number of repetitions irrespective of the number of post-synaptic spikes, whether they are singlets (Fig. 5A) or doublets (Fig. 5C). The nature of the synaptic rule presented here tells us that on singlet input-output pairings, the somato-dendritic coupling boosting is not enough to activate HVA-Ca^2+^ channels that would increase Ca^2+^ levels when single spikes occur. This implies that the present model lacks plasticity mechanisms for the induction of strong LTD at low stimulation frequencies. Nevertheless, strong LTD can still occur when the plasticity signals relax after sufficiently many bursts of spikes (Supplementary Fig. 3), which homeostatically regulate synaptic conductance changes.

### Learning reward-place association on behavioral timescales

As shown above, our learning rule yields a Mexican-hat-like STDP time window consistent with experimental observations. However, a well-recognized issue about STDP is that it only describes synaptic weight changes on a relatively short timescale of the millisecond range^27^. Then, a question arises about whether our learning rule can associate neuronal events on short timescales with behavioral events on longer timescales. We address this question in a reward-place association task, a typical task for studying behavioral timescale plasticity (BTSP)^30^.

When rodents are tasked to walk or run along a treadmill to find a given reward location, CA1 pyramidal cells rapidly adapt their spiking activity to encode place locations in the new environment. In this case, place fields would over-represent the spatial location associated with reward-related information. This facilitation may be explained by a dual association of reward and place information by single CA1 neurons.

As NMDAR blockade is known to impair the formation of place cells and reward location associations^51^, we exploited the place-encoding capacity of the Ca^2+^/NMDAR-based model and study the effects of NMDAR blockade in this task. We ask whether BTSP can emerge from the slow activation-deactivation dynamics of NMDARs and the calcium influx through these receptors.

In our modeling of this task, the dendritic compartment receives place information and reward-related information (mimicking inputs from hippocampal CA3 and the ventral tegmental area VTA, respectively). ‘Reward’ is delivered at a fixed location between the two endpoints of the treadmill (Fig. 6A and Supplementary Fig. 4). Other details of the simulation setting are described in the Methods. To properly simulate the behavioral paradigm, we constructed a finite state machine (FSM) that simulates the realistic behavior of a rodent running along a treadmill. This agent integrates the animal’s position with respect to time given their acceleration and velocity as they traverse five different behavioral states during exploration, namely, “accelerating”, “peak velocity run”, “decelerating”, “reward (licking)”, and “no reward”. For example, the “decelerating” state is a transitory state that mimics sudden stopping with reward proximity. See Supplementary Fig. 5 for details of the FSM. As frequently observed in rodent behavior, we mimic rodent licking in unrewarded and rewarded areas by reducing their acceleration towards a minimal speed along the treadmill for a given time sampled from a uniform distribution. This enhances spiking activity received at specific times (red colored spans on Fig. 6B).

**Figure 6:**
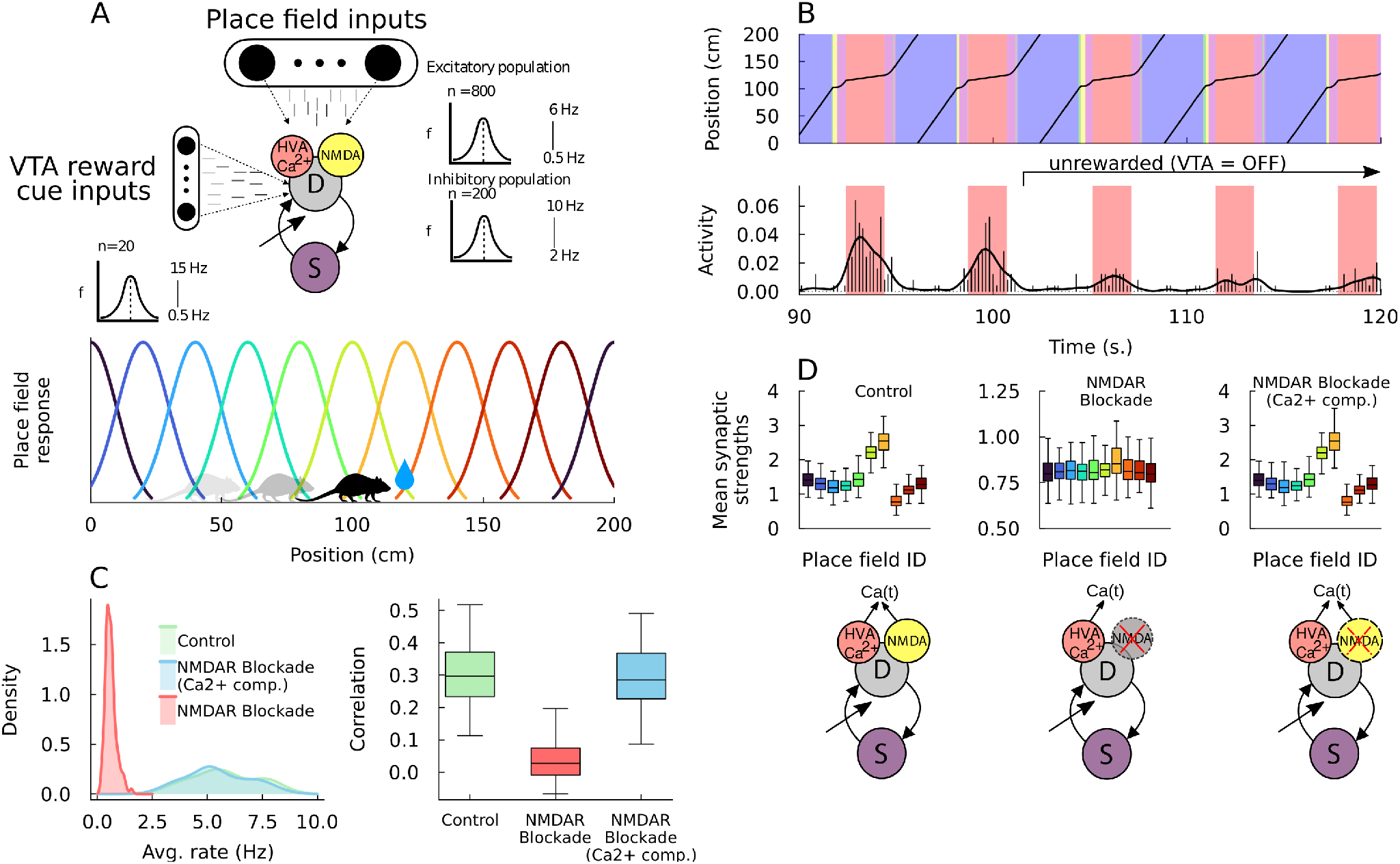
Reward-place associations by the Ca^2+^/NMDAR-based neuron model. (**A**) A Ca^2+^/NMDAR-based neuron model receives place information and reward information from separate groups of Poisson spiking input units, which may correspond to the hippocampal area CA3 and VTA, respectively. The sizes of these neural populations and their peak and minimum firing rates are shown on inset plots. The positions of CA3 place fields along the treadmill are highlighted in colors and the reward location is marked by a water drop (bottom). (**B**) Top, Five behavioral states of the FSM are indicated by their positions along the track during an example trial: “accelerating” (pink); “peak velocity run” (blue); decelerating (green); reward licking (red); no reward (yellow). Bottom, The histogram of neural activity (in %) is shown for the example trial. Colored spans denote `REWARD` state but reward cue inputs are silenced after the 100-second mark. Neural responses during the unrewarded period (100-second mark onwards) are used for calculating performance metrics. (**C**) Left: the distributions of trial-averaged output spike rates for a reward-off period (from 100 to 180 seconds in each trial) are shown for the three model conditions. Right: box plot of correlations between output spikes and reward period are shown for the three model conditions. (**D**) Box plot of mean synaptic strength of place field inputs for all model conditions (diagrams below). Colors as in the place-field inputs described in (**A**). All box plots in this figure contain five-number summary measures without outliers.

We test our model in the following three different conditions for a total of 100 trials per condition (Fig. 6C). In condition 1, we use the Ca^2+^/NMDAR-based model with no changes to test the capacity of the model to associate reward and place information. In the condition 2 (NMDAR blockade condition), we block NMDAR dynamics by invalidating their equations (i.e., we set *I*_*NMDA*_ = 0 and *I*_*Ca*_ = 0 in Eq. 30) and test the capacity of the model. Mathematically, this NMDAR blockade makes the model equivalent to the model that only includes HVA-Ca^2+^ channels presented in Fig. 2C. Finally, in condition 3 (Ca^2+^ comp. condition), we compensate for calcium intrusion via the NMDAR channel to study the effect of its slow calcium dynamics. This compensation occurs by enabling the fraction of Ca^2+^ current influx through NMDAR into the dendrite while blocking the voltage influences of the NMDA current (i.e., we set *I*_*NMDA*_ = 0 while leaving *I*_*Ca*_ intact in Eq. 30). This fraction represents around 5% (parameter *f* in Eq. 30) of the total NMDA current but significantly affects the model’s performance in learning, as demonstrated below.

In Fig. 6D, we assess the performance of the model by computing the correlation between spiking activity and a binary reward signal, which takes the value 1 during the “reward” state (i.e., red-shaded spans on Figure 6B) and the value 0 otherwise. Our Ca^2+^/NMDAR-based model achieves a reasonably good performance in condition 1, suggesting that the reward-place association has been learnt when NMDAR is intact. A perfect correlation should not be expected, as spiking activity can occur outside the reward state and may not occur during the whole reward state. In contrast to condition 1, the model is unable to learn this association in the NMDAR blockade condition. Interestingly, however, learning capability is recovered when compensating for the Ca^2+^ influx through NMDAR dynamics (Ca^2+^ comp. condition). We note that the firing rate distributions of cell models in the control and Ca^2+^ comp. condition are overlapping significantly whereas, in condition 2, the blockade of NMDAR heavily affects spiking activities. We further clarify performance differences between the different conditions by plotting the average values of synaptic strengths belonging to each place field across all trials (Fig. 6D). The results show that the NMDAR blockade greatly disrupts plastic changes of synapses necessary for place-reward association.

Altogether, our results suggest that the slow rising/decaying dynamics of NMDAR-mediated calcium influx are necessary and sufficient for associating neuronal spiking activity occurring on short timescales with behavioral events (reward delivery) occurring on longer timescales. Importantly, the calcium current through the NMDARs represents only a small fraction of the total NMDA current. Nevertheless, when subjected to unstructured high-frequency inputs, the proposed Ca^2+^/NMDAR-based learning rule can associate the unstructured inputs to a neuron model with behavioral information streams arriving at longer timescales. Thus, our model describes a biologically plausible mechanism underlying behavioral timescale plasticity.

### Few-shot learning by pre-existing cell assemblies

Now, we investigate whether pre-existing assemblies of the proposed neuron model improve the detection of temporal communities in afferent inputs. For this purpose, we constructed a recurrent network of 400 excitatory and 100 inhibitory units with 20 pre-configured excitatory cell assemblies with average size of about 18 units each. To reduce the load of numerical simulations, we used the spike trace-based two-compartment model defined in (Eq. 1). For a comparison reason, we used the same input scheme as used in Fig. 1A), which contained three repeated patterns. Connections from excitatory and inhibitory afferent input units to network units are all-to-all, with Gaussian-distributed synaptic weights (Fig. 7A). All excitatory and inhibitory connections in the network model were modifiable according to the proposed learning rule. We evaluated the ability of the recurrent network to detect any of the salient patterns amidst noise. Network parameters used are presented in Table 4.

**Figure 7:**
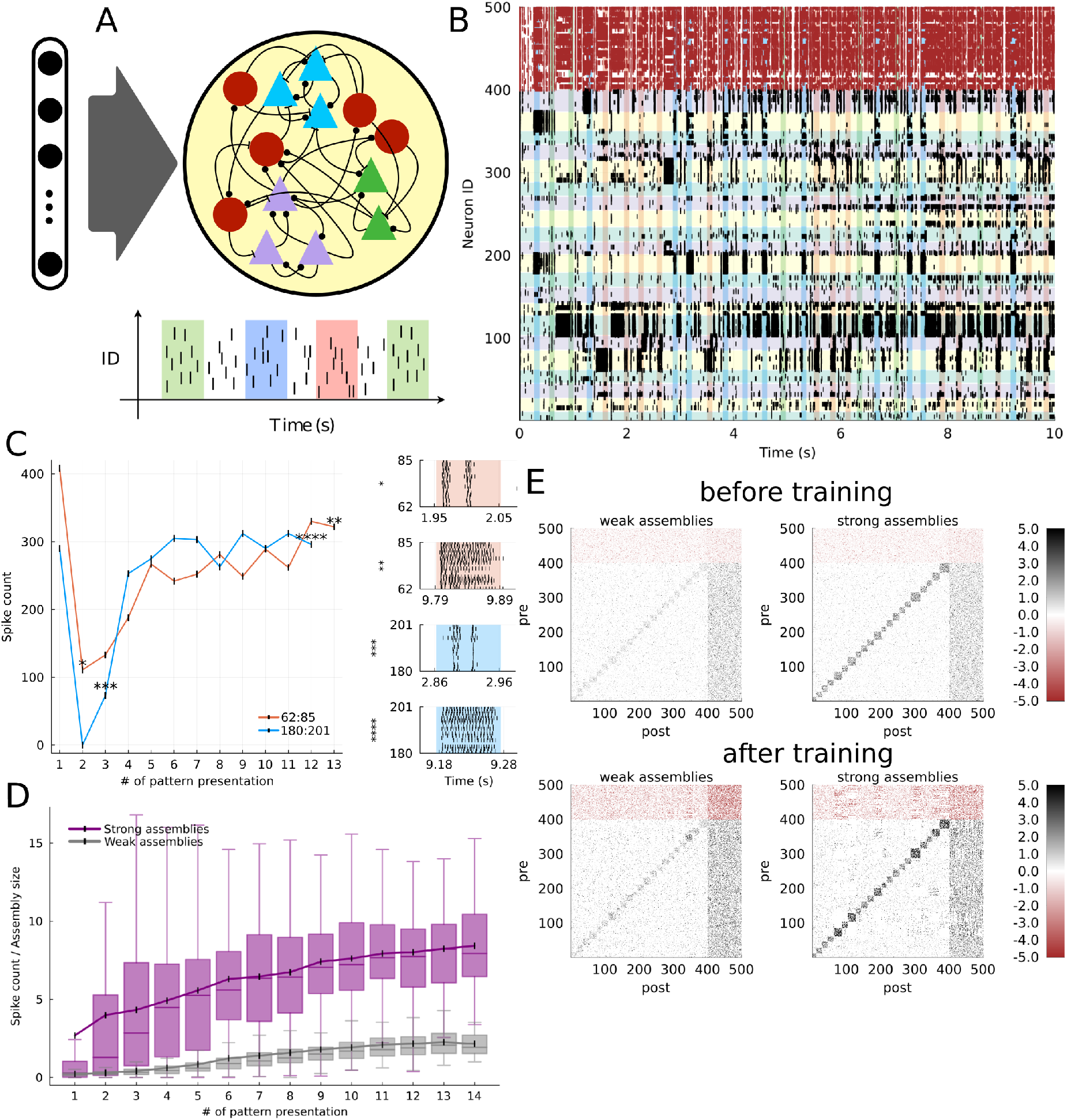
Few-shot learning through competition among pre-existing assemblies. (**A**) Recurrent network with pre-existing cell assemblies receive all-to-all afferent input from an input layer used in Fig. 1A. (**B**) Raster plot of network activity in a 10-second trial. Neurons 1 to 400 are excitatory (black) and 401 to 500 are inhibitory (redbrown). Presentations of three patterns are indicated as vertical spans and pre-existing assemblies as horizontal spans. (**C**) Left, Spike counts of two cell assemblies are plotted for a given number of pattern presentations. The asterisks refer to the corresponding right panels. Right, Raster plots of assembly activity for given pattern presentation and given time indicated in the left panel. (**D**) Box plots with five-number summary measures (outliers excluded) and mean value of tuned assemblies’ spike counts relative to assembly size are shown for strongly (purple) and weakly (gray) coupled initial assemblies (20 trials each) across presentations. (**E**) Heatmaps of synaptic weights before (top) and after training (bottom) for trial in B truncated between -5 and +5 for visualization.

With appropriate network connectivity and initial synaptic weight distributions, the network self-organized robust response patterns in a competitive manner, wherein neurons belonging to a pre-existing assembly were selectively co-activated by one of the salient input patterns. In Fig. 7B, cell assemblies could learn such selective responses at the early stage of learning as long as their within-assembly connections were strong enough. Due to the nature of the recurrent connectivity, it was rarely the case that a single assembly of neurons grew up to dominate the activity of the entire network. For instance, a highly active assembly of neuron 104 to neuron 127, which became selective to none of the patterns, neither impeded a strong response to pattern red of the assembly of neuron 62 to 85 nor the response to pattern blue of the assembly of neuron 180 to 201. To access the rapid learning by pre-existing cell assemblies quantitatively, we show the spike count occurring during each pattern presentation across the 10-second simulation for the example trial (Fig. 7C, left). To our surprise, very few presentations were sufficient to develop robust responses to salient patterns. The growth of neuronal responses is also indicated by the responses of cell assemblies 62-85 and 180-201 during the early and late phases of pattern learning (Fig. 7C, right). However, some pre-existing cell assemblies could learn no selective responses to patterns, as exemplified by the assembly 104-127 in Fig. 7B. Because the network could not learn robust responses without the plasticity of inhibitory neurons, inhibitory plasticity is likely to be crucial for the stability of pre-existing cell assemblies and the formation of their pattern-selective responses.

While few-shot learning of patterns is remarkable, it is unclear whether strongly coupled assemblies are necessary for such a fast convergence. To address this, we run a set of 20 networks endowed with the same connectivity in two different conditions (i.e., 40 trial simulations in total). In a scenario, excitatory assemblies are strongly connected (as is the case in Fig. 7B) with Gaussian distributed weights with the mean *μ* = 3.0 and standard deviation *σ* = 1.0, while in a second scenario, within-assembly synaptic weights are weak and defined with Gaussian parameters *μ* = 0.0 and *σ* = 1.0 (the weights are always positive). The spike counts normalized by assembly size are plotted against the number of pattern (any) presentations in Fig. 7D. Overall, the strongly coupled pre-existing cell assemblies are capable of producing, on average, a robust spiking response from the very beginning of the simulation whereas the weakly coupled ones require many more presentations. In terms of time-to-convergence, the number of presentations required for weakly coupled assemblies generally matches that of the single-cell model presented in previous figures. The ability of pre-existing cell assemblies to develop a robust spiking response to statistically salient patterns is further enhanced by the strengthening of with-in assembly excitatory connections. As indicated by the heatmaps of the recurrent weight matrix before and after training in each condition (Fig. 7E), such strengthening occurs in strongly-coupled pre-existing assemblies but not in weakly-coupled ones. We also observe that cell assemblies become tightly coupled with the recurrent inhibition that enforces network stability.

In summary, we found that the proposed learning rule implemented at afferent and recurrent synapses enables few-shot unsupervised learning of temporal communities of presynaptic neurons. Pre-existing cell assemblies embedded in a recurrent neuronal network further accelerate this pattern detection. Importantly, the plasticity of inhibitory synapses, which we assumed obey the same proposed learning rule, is necessary for stabilizing network responses to patterned synaptic inputs.

## Discussion

In this study, we have proposed a synaptic plasticity rule for cortical pyramidal neurons and implemented it in a two-compartment neuron model. The learning rule is inspired by machine-learningbased predictive learning for single cortical neurons and supported by findings in cortical neurobiology. Like a machine-learning counterpart, the two-compartment model is endowed with computational features to detect and robustly spike to hidden patterns amidst noisy input. Our model predicts that a transient boosting of the somato-dendritic coupling is crucial for the pattern learning ability and NMDARs significantly improve the SNR of this learning. The proposed learning rule can replicate the STDP rule observed for spike bursts at the hippocampal Schaffer collateral. Furthermore, pre-existing cell assemblies greatly accelerate pattern learning in a recurrent network of two-compartmental neurons.

### A biologically plausible single-cell mechanism to learn activity patterns

The capacity of PCs to effectively adapt their neuronal responses to stimuli is largely mysterious. Various plasticity rules have been proposed to understand how PCs assign credit to the relevant synapses. Additive STDP leads to efficient close-to-optimal pattern detection on single neuron modeling^9^. Voltage-based STDP rule supports pattern detection in models with plausible neural circuitry, and dendritic voltage-based STDP preserves learned associations across longer periods of time^10^. STDP is particularly useful in competitive neural networks^13,39^. While STDP rules may be a simplistic abstraction of the complex biological processes that underlie synaptic plasticity events, machine learning-based approaches and the resultant nonlinear Hebbian rules aim at optimal performance on complex tasks.^6,8,11,12^. Our work based on unsupervised predictive learning sits among this class of algorithms while offering a biologically plausible realization, thus bridging the two classes of synaptic plasticity rules.

Specifically, we highlight that our biological learning rule does not require an extensive presentation of patterns to credit the relevant synapses rapidly. To achieve this rapid learning, our model hypothesized a spike-triggered transient boosting in somato-dendritic coupling, which enables bAPs to propagate into the dendritic compartment. Cortical PCs exhibit a high correlation between somatic and dendritic activities, and bAPs are suggested to underlie associative plasticity^27,52^. Although this transient boosting needs to be confirmed by experiment, the somata and dendrites of *in-vivo* L5 PCs display widespread coupling asymmetry and high correlation of Ca^2+^ transients especially during somatic burst spiking^53–55^. In our model, the transient coupling boosting allows the invasion of bAPs into the dendritic compartment only during neuronal refractoriness, forming a positive feedback loop between somatic firing events and dendritic Ca^2+^ plasticity updates to improve the SNR.

Transient boosting in dendritic coupling is thought to depend on various ionic channels that support the mechanistic transmission and regeneration of bAPs. Specifically, Na+ and K+ channels and their associated densities across the dendritic branches may underlie bAP transport to distal dendritic branches^52,56,57^. In principle, we may experimentally test the transient change of the active component of the coupling conductance by looking at the partial derivative of the membrane resistance along the proximal dendrites of PCs. However, measuring a rapid change in the axial resistance and its effect on the transitory part of signal transmission is technically difficult.

### Underlying molecular mechanisms of our calcium-based learning rule

Our error signal 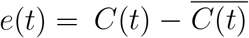 (Eq. 5), which measures the necessary amount of plasticity induction, depends on the long-term average of the intracellular calcium concentrations. Although we did not explicitly model this averaging process, it may involve interactions between free Ca^2+^ ions and calmodulindependent protein kinases such as CaMKII and CaN, which are known to affect LTP and LTD, respectively^58–60^ (see Yasuda et al.^61^ for a recent review). In particular, CaMKII likely acts as a leaky integrator of Ca^2+^ during LTP induction but not during maintenance of synapses^62^. Our model employed the multiplicative bounding function (Eq. 15) to maintain the learned synaptic representations by suppressing a further LTP induction. Importantly, CaMKII binding rates are fast and this kinase acts as soon as Ca^2+^ ions flow into the dendritic spine^63^. We conveniently chose an appropriate decay constant for 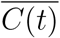 that was sufficient for the task, but whether this constant matches the decay constant of intracellular kinase messaging remains to be understood.

The spike-trace-based plasticity rule relies on the artificially generated graded spike trace *Y* (*t*), but biological cells do not have a readily available trace that informs spiking activity. With this and the vast amount of existing literature on calcium-based synaptic plasticity in mind (e.g., Inglebert et al.^44^), we replaced the artificial trace with the calcium concentration trace *C*(*t*) in our calcium-based plasticity rule. Thus obtained calcium-based plasticity rule yielded a symmetric spike-timing rule similar to the Shaffer collateral STDP for spike singlet-doublet pairings. This contrasts with the predictions of a previous calcium-based STDP model for spike singlets, in which different levels of calcium entry result in stages of LTD after a given Ca^2+^ concentration threshold and those of LTP after at higher levels of Ca^2+^ concentration^64^. It is noted that the Shaffer collateral also exhibits a symmetric STDP under dopaminergic modulation^65^. We consider our plasticity rule to be a generalized account of the course of synaptic plasticity signals with no strict requirements regarding threshold constraints or LTD/LTP amplitude specifications.

### Burst-triggered symmetric STDP in the calcium-based model

The proposed calcium-based plasticity rule produces synaptic strength changes resembling those observed in the burst-triggered STDP at the CA3-CA1 Schaffer collateral^26,50^. The results are interesting as bursts are thought to transmit spikes more faithfully than single spikes^15,66,67^. On the other hand, our model (and the experiment as well^50^: see Fig. 5A) did not produce an asymmetric STDP time window reported in other studies for singlet-spike paring at the Shaffer collateral^23,24^. It was speculated that this discrepancy was partly due to differences in an experimental setting^50^: while a potassium-based solution induces LTD and hence gives asymmetric STDP, a cesium-based intracellular solution blocks potassium channels and depolarizes the postsynaptic neuron to enhance bAPs, which may turn LTD into LTP to generate symmetric STDP^24^.

The Schaffer-collateral STDP only exhibits LTP for a small number of pairings (see Fig. 5B). However, for a large number of singlet-doublet pairings, strong LTD components appear at both sides of the LTP time window (Fig. 5C). The Schaffer-collateral STDP is also LTD dominant in singlet-singlet pairings (Fig. 5A). Our neuron model claims that the LTP components for single-doublet pairing arise from a reliable bAP propagation (i.e., a sufficient invasion into dendritic branches). However, our model cannot account for the LTD components. A likely reason for the lack of LTD is that our STDP assessment ignores the presence of inhibition, which was suggested to induce LTD for a causal pre-post spike pairing^24,25^. In addition, our model does not include Ca^2+^-dependent K^+^ channels, which are known to halt positive loops mediated by Ca^2+^ signaling^68–70^. These channels suppress the activation of CaMKII in spines^71^, a kinase crucial for the LTP induction, and decrease intraburst firing rates, which may induce LTD^72^. All these possibilities have yet to be addressed experimentally.

### Emergence of BTSP in the Ca^2+^/NMDAR-based plasticity model

In our model, NMDAR-mediated Ca^2+^ influx significantly increases the SNR of neuronal responses to patterns responses (Fig. 3C). NMDAR is heavily implicated in LTP, and its role in place field development and retention during novel environment exploration is also established^51,73^. Our model suggests that NMDAR-mediated Ca^2+^ influx is necessary for associating reward and place information during the exploration of novel environments (Fig. 6A and Suppl. Fig. 5). The result is consistent with the fact that the blockade of NMDARs significantly impairs place field formation^74^. It is worth noting that this reward-place association requires a plasticity rule that extends over a behavioral timescale of seconds^30^. Our model achieves this seconds-long association by a small fraction of Ca^2+^ influx through NMDARs (Fig. 6D). A mathematical formalism of BTSP involves an instructive signal triggered by dendritic plateau potentials^75^. In contrast, our model suggests that a bAP-triggered transient boosting of somato-dendritic coupling conductance and Ca^2+^ influx through NMDARs bestow the observed rapidity and high SNR to the seconds-long associative plasticity.

### The role of pre-existing cell assemblies in rapid pattern detection

The ability of the brain to rapidly learn and remember features of sensory experiences has long been thought to require the experience-dependent generation of cell assemblies. However, whether and to what extent the learning of cognitive experiences is constrained by pre-existing cell assemblies in the brain has been debated^32–34,76^. Our network modeling results showed that the proposed learning rule rapidly associates pre-existing assemblies activated repeatedly by patterned inputs with one of the patterns when within-assembly connections are strong enough (Figs. 7C, D). The stability of network responses is maintained during this rapid association by reorganizing recurrent inhibition (Fig. 7E). These results suggest that strongly coupled pre-existing assemblies that compete for stimuli provide cortical machinery for rapid learning of activity patterns. In machine learning-based approaches, competition among neurons during training can prevent overfitting by encouraging simpler and more general representations^77^. In the brain, assemblies of tightly connected neurons can compete for simple and robust representations of information carried by stimuli. As the identity of the assembly may change over time, which neuronal assembly gets recruited and wins over the rest may not matter^78,79^.

### Limitation of the present model

The dendrite of our neuron model likely describes basal dendritic branches receiving sensory input rather than apical dendritic branches receiving top-down input. The nature of our Hebbian learning rule relies on strong somato-dendritic correlations during spiking activity, which is reminiscent of strong correlations between basal dendritic Ca^2+^ events and somatic firing events in PCs. In contrast, evidence indicates that the correlation between Ca^2+^ events and somatic spikes are weak in apical tuft dendrites^43,80^ (but see^53^). Indeed, the strengthening of synapses highly depends on dendritic location^2,81,82^. Then, the question arises about how cortical neurons integrate bottom-up sensory information and top-down context information arriving at different dendritic locations^83,84^. The present model falls short for addressing this question. Exploring such integration requires more realistic separate compartmentalization^15,54,55^ combined with compartment-specific plasticity mechanisms^2,82,85^.

Our simplified models did not incorporate several ionic channels that can contribute to synaptic plasticity. With rising levels of Ca^2+^, SK channels are known to terminate NMDAR activation via decreasing the local potential^69^. These channels may exert a significant effect on BTSP, and a future model needs to address this effect in a realistic morphological model with localized spines and intracellular Ca^2+^ pools. Given the non-linearity of intracellular processes, we suspect that an extensive remodeling work is necessary to translate the Ca^2+^ transients into localized compartments across morphologically realistic dendritic branches. For instance, appropriate channel densities might be required to address bAP invasion in the dendritic tree and subsequent synaptic plasticity cascading in its branches ^40,41^. A faithful propagation of bAPs is crucial for “self-supervising” the unsupervised learning proposed in this study.

In sum, we have modeled the neurobiological mechanism underlying a rapid adaptation of pyramidal cell spiking responses to a statistically salient pattern hidden in the barrage of synaptic input. Our model predicts the elements of single-cell and network-level computations, including a transiently boosted somato-dendritic coupling and pre-existing cell assemblies, crucial for this learning. The proposed mechanism is consistent with the behavioral time scale learning and may underlie one-shot learning of episodes, a great advantage of the brain’s memory systems.

## Acknowledgement

This work was partly supported by JSPS KAKENHI no. JP23H05476. The authors are grateful to Yukiko Goda and Bernd Kuhn for the valuable discussion on the physiological properties of pyramidal neurons. We are also grateful to Milena M. Carvalho and Toshitake Asabuki for their helpful discussions and support.

## Data availability

All numerical datasets necessary to replicate our results can be generated with the software code referenced below. No datasets were generated during this study.

## Code availability

All codes were written using standard libraries with Julia programming language version 1.8 and some visualizations in Python 3. Example program codes used for numerical simulations and data analysis are available at https://github.com/gsivori/transientboost/.

## Methods

The following sections describe model equations in separate subsections. All parameter values discussed in these subsections are shown in Table 1, 2, and 3 for presented models.

**Table 3:**
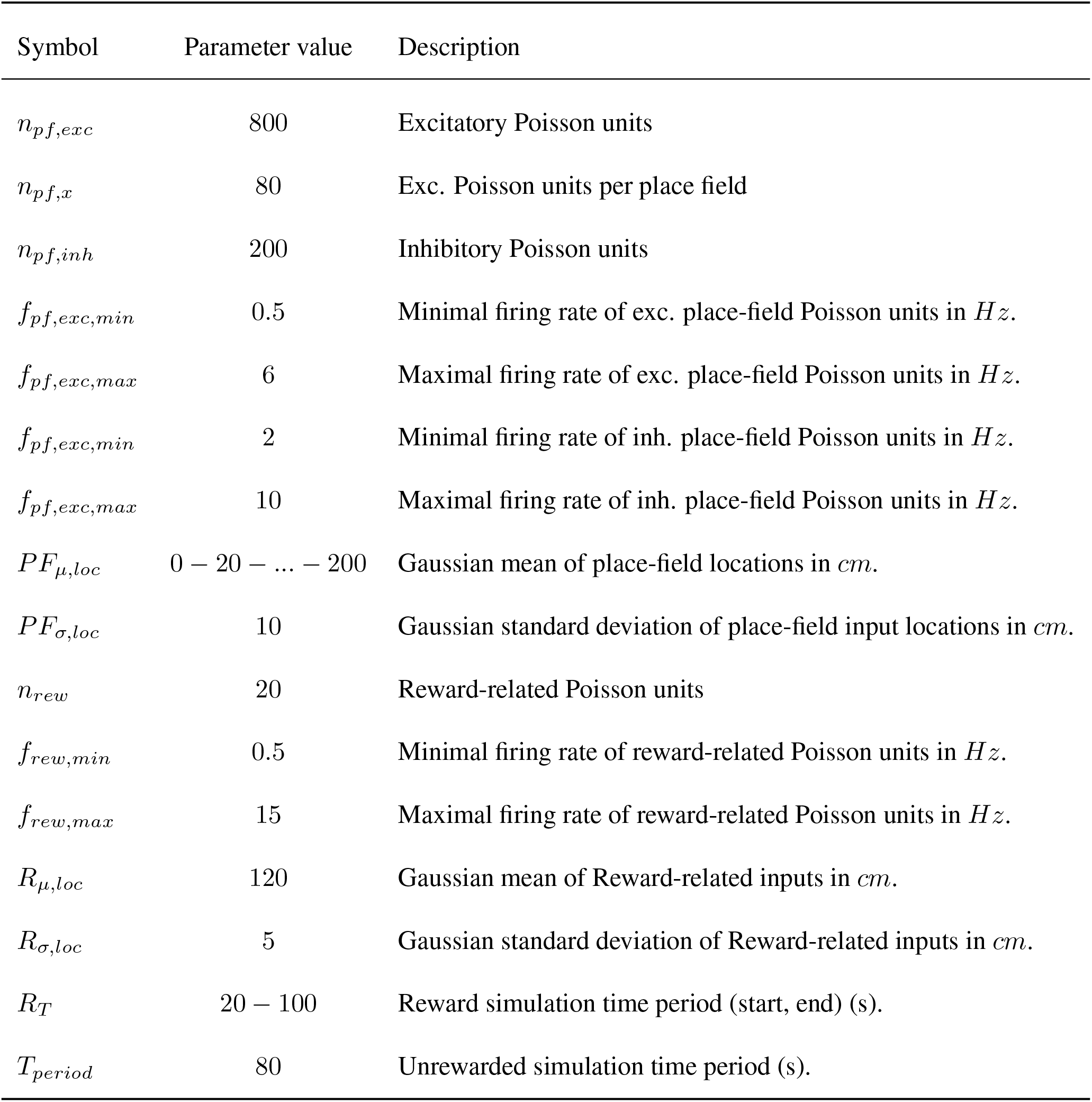
Reward-place association task parameter symbols, values, and descriptions.

**Table 4:** Recurrent network with assemblies parameter symbols, values, and descriptions. Most cell parameters as in Spike trace-based model from Table 1 unless otherwise noted.

| Symbol | Parameter value | Description |
| --- | --- | --- |
| $C_x$ | exc=180,inh=150 | Somatic membrane capacitance in $pF$ |
| $N_x$ | exc=400,inh=100 | Number of excitatory and inhibitory recurrent units |
| $p_{e \rightarrow e}$ | 0.1 | Connection probability from Exc.-Exc. units |
| $p_{e \rightarrow i}$ | 0.3 | Connection probability from Exc.-Inh. units |
| $p_{i \rightarrow e}$ | 0.4 | Connection probability from Inh.-Exc. units |
| $p_{i \rightarrow i}$ | 0.5 | Connection probability from Inh.-Inh. units |
| $W_{e \rightarrow e}$ | LogNormal($-0.42, 0.79$) | Exc.-Exc. synaptic weight distribution (mean,std) |
| $W_{e \rightarrow i}$ | LogNormal($0.48, 0.63$) | Exc.-Inh. synaptic weight distribution (mean,std) |
| $W_{i \rightarrow e}$ | LogNormal($-0.74, 0.96$) | Inh.-Exc. synaptic weight distribution (mean,std) |
| $W_{i \rightarrow i}$ | LogNormal($-0.39, 0.78$) | Inh.-Inh. synaptic weight distribution (mean,std) |
| $p_{cl}$ | 0.5 | Connection probability of assembly units |
| $n_{cl}$ | Normal($18.0, 3.0$) | Assembly size (integer) distribution (mean,std) |
| Strong $W_{cl}$ | Normal($3.0, 1.0$) | Strong assembly synaptic weight distribution(mean,std) |
| Weak $W_{cl}$ | Normal($0.0, 1.0$) | Weak assembly synaptic weight distribution (mean,std) |
| $\eta_{aff}$ | 0.05 | Learning rate for afferent inputs |
| $\eta_{rec}$ | 0.05 | Learning rate for recurrent inputs |

### Somatic compartment (spike trace-based)

The somatic membrane potential *V*_*s*_ integrates input coming from the dendrite through the conductance *g*_*cds*_ and leaks out through the conductance *g*_*ls*_. *V*_*ls*_ represents the leakage potential of soma, *C*_*s*_ is the capacitance of the somatic compartment, and *V*_*d*_ represents the dendritic membrane potential. The potential evolves following equation 6:

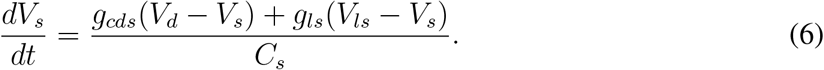

When *V*_*s*_ *> V*_*th*_, a spike event occurs and the somatic membrane potential is updated to a reset voltage *V*_*s*_ *→ V*_*re*_ where it remains refractory for *τ*_*ref*_ between 2 to 3 milliseconds (in all models). During a refractory period, the integration of equation 6 is halted, and resumed afterward.

### Dendritic compartment (spike trace-based)

The dendritic membrane potential *V*_*d*_ receives excitatory and inhibitory input through conductance-based currents. With *g*_*e*_ and *g*_*i*_ the excitatory and inhibitory conductances, and *V*_*er*_ and *V*_*ir*_ the excitatory and inhibitory reversal potentials, the dendritic potential evolves following equation 7:

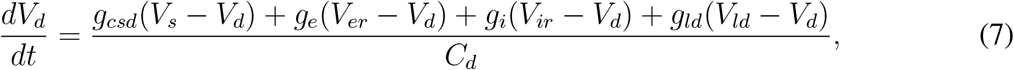

where *g*_*csd*_ is the conductance parameter for somatic to dendrite compartment, *g*_*ld*_ is the leakage conductance of the dendrite, *V*_*ld*_ the leakage reversal potential of the dendrite, and *C*_*d*_ is the capacitance of the dendritic compartment.

We model the excitatory and inhibitory conductances as double exponential functions with appropriate time rise and time decay constants. These are updated using equations 8 to 11:

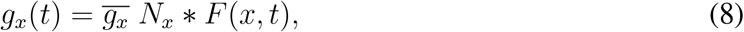

with *x* being excitatory or inhibitory, 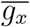 the peak conductance value, *N*_*x*_ a normalizing constant, and *F* (*x, t*) a double exponential function that evolves according to the following equations 9-11:

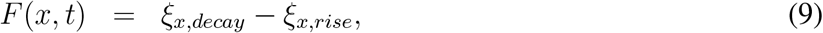

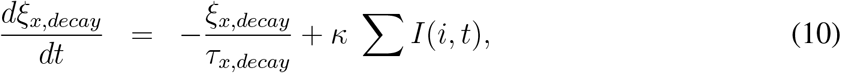

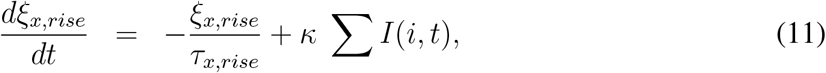

with *I*(*i, t*) = *w*(*i*)*∗S*(*i, t*) the synaptic input *i* at time *t. S*(*i, t*) the binary input matrix representing the timing of spikes. And *κ* is a homeostatic balancing parameter (see below). *N*_*x*_ is calculated using equation 12:

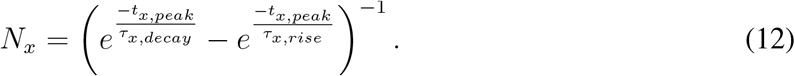

Finally, the time of the peak *t*_*x,peak*_ is obtained by equation 13:

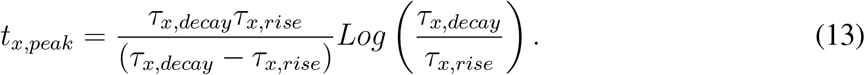

### Spike trace-based plasticity rule

Our proposal is a multiplicative Hebbian synaptic plasticity rule. In Eq. 1, the postsynaptic potential term consists of the magnitude of the afferent input *w*(*i*) that decays over time with a time scale activation *τ*_*p*_:

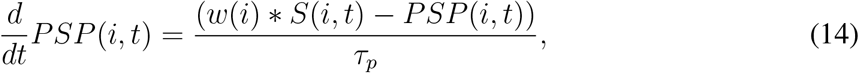

with *S*(*i, t*) the binary input matrix representing the timing of spikes.

The last term in Eq. 1 consists of an Alpha function with free parameters *ϕ*_*ζ*_ and *τ*_*ζ*_ chosen to accommodate the growth of synapses without requiring explicit boundaries.

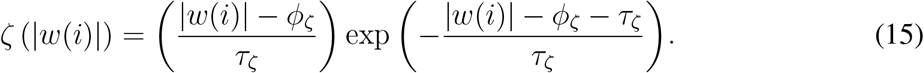

At any given time, *PI*(*i, t*) corresponds to a sample of inducible plasticity for a given synapse *i*. Synaptic weights are gradually evolved over time by sampling from this trace (Eq. 16) at a learning rate *η*:

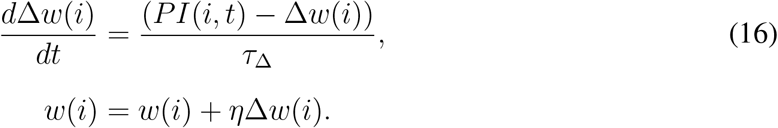

Finally, we avoid plastic changes that change the nature of the synapse (excitatory to inhibitory and vice-versa) following equation 17 (Dale’s Law):

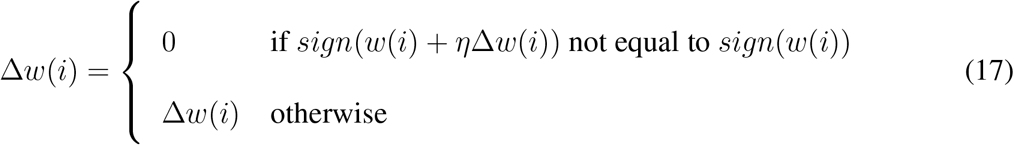

### Somatic compartment (calcium-based)

The somatic compartment for our extended model (Calcium-based) requires the upswing potential *V*_*peak*_ after the spike event (i.e. when *V*_*s*_ *> V*_*th*_). To mimic the quick rise and decay of a biophysical action potential, we chose the values of leak currents during and outside the refractory period such that *g*_*lsr*_ *>> g*_*ls*_. The somatic membrane potential follows equation 18 outside the refractory period, and equation 19 during the refractory period

where the leakage conductance changes to *g*_*lsr*_:

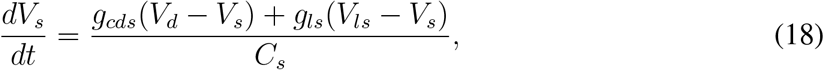

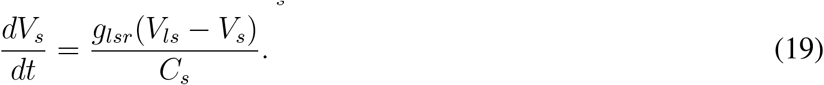

### Dendritic compartment (calcium-based)

We include the calcium current description based on the dynamical rates obtained with Hodgkin-Huxley steady-state activation variables and forwardbackward rate constants (*α* and *β* respectively) for HVA-Ca^2+^ current^42^. The rate constants depend on dendritic voltage *V*_*d*_ and are measured at all time steps following the sets shown in equations 20-21.

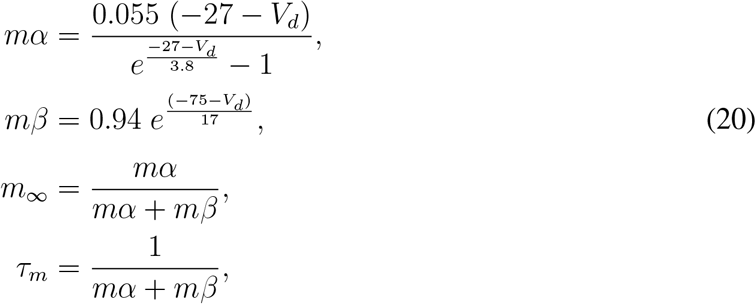

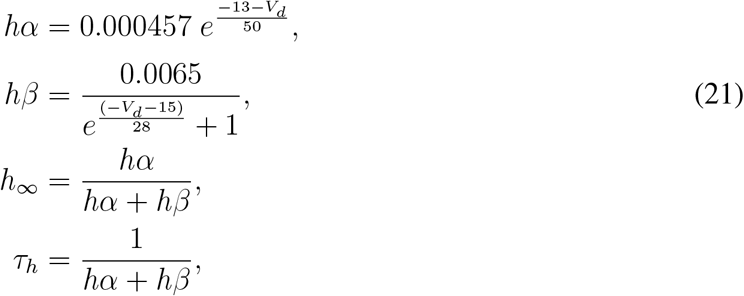

with parameters *m*_*∞*_, *h*_*∞*_, *τ*_*m*_, and *τ*_*h*_ defined dynamically, the activation and deactivation variables are integrated over time using equations 22 and 23, respectively.

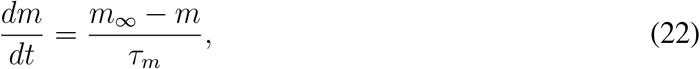

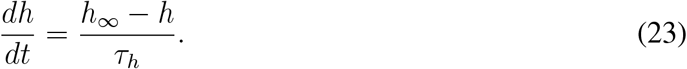

Following the Ca^2+^ current description in^42^, we model this current with *m*^2^*h* kinetics as shown in equation 24. In this equation, *g*_*Ca*_ is the maximum calcium current conductance, and *V*_*Ca*_ is the calcium reversal potential. Note that the defined direction of the calcium is inwards, i.e., depolarizing the compartment.

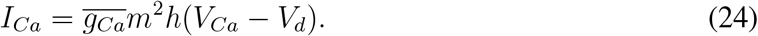

Thus, the dendritic compartment evolves its membrane potential using the following equation 25. *I*_*csd*_ is the somato-dendritic coupling current which will be discussed below:

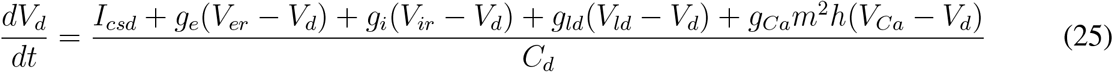

### Somato-dendritic coupling current (calcium-based)

Our description of the transient change of the somato-dendritic coupling *g*_*csd*_ relies on the biophysical aspects that occur during BAC firing. To start with, in biological cells, conductance between dendrites and the soma of the cell have both active and passive properties, and the active properties rely on the locality of the channels that are presently opened or closed. Since inputs that traverse the dendritic branches during non-refractory states propagate towards the soma, the coupling conductance is fairly constant dominated by its passive properties. On the other hand, when the neuron is engaged in firing an action potential –and thus entering the refractory state– this coupling is dominated by active properties as the soma experiences a diverse opening and closing of channels that repolarize the cell.

We model this behavior by transiently affecting the somato-dendritic coupling current during refractory as is shown in equation 26:

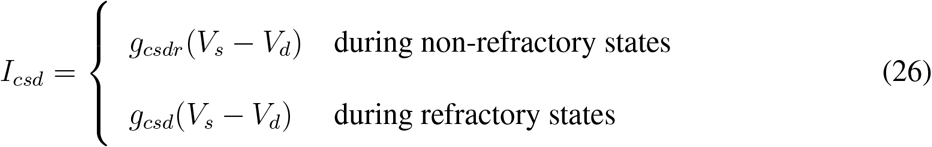

The transient increase of the coupling *g*_*csd*_ occurs on every spike. Similarly to *g*_*e*_ and *g*_*i*_, the coupling conductance evolves over time following the equations 8-13. The parameters for this coupling are *τ*_*csd,rise*_, and *τ*_*csd,decay*_ for the rise and decay of the double exponential function. *t*_*peak,csd*_, and *N*_*csd*_ as normalizing constants. *g*_*csdr*_ as the resting coupling value, and *m*_*csd*_ as the amplitude of the increase in nS. On each spike, the exponentials *ξ*_*csd,decay*_ and *ξ*_*csd,rise*_ receive *m*_*csd*_ as input (bAP strength/pulse) at *t* = *t*_*f*_, where *t*_*f*_ is the somatic spike time. The set of equations 27-29 describe the transient increase of coupling *g*_*csd*_.

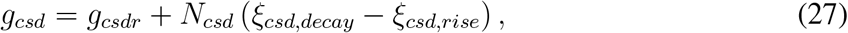

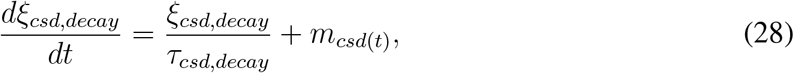

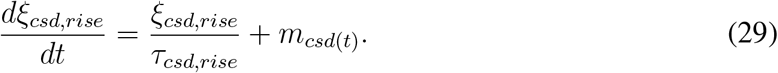

### NMDAR dynamics (Ca^2+^/NMDAR-based)

Our last model includes NMDAR dynamics based on the set of equations 8 to 11 and a conductance-based current obtained from the following:

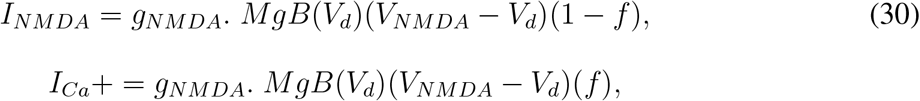

with *f* a constant factor mediating NMDA-dependent calcium transient influx *I* _*Ca*_. Magnesiumblock voltage-dependence function was taken from Jahr & Stevens (1990)^48^:

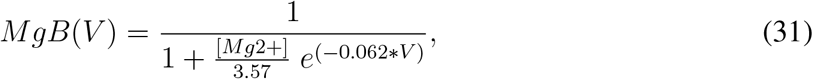

with Magnesium concentration [*Mg*2+] in *mM* . Time constant parameters for NMDAR are originally temperature-corrected values (Q10). Importantly, we consider that around 75% of the excitatory synapses in the model also activate NMDAR dynamics. This input is thus affected by integrating *g*_*NMDA*_ as other conductances (see Eqs. 8-13).

### Homeostatic plasticity

In all presented models, we introduce homeostatic synaptic plasticity to keep the overall synaptic structure of inputs balanced across the simulation. This is done simply by updating a synaptic constant that multiplies all synapses at the moment of their integration using equations 10-11. Thus, on each time step, synaptic input is multiplied by *κ*. We update this constant every *T*_*ϑ*_ milliseconds by preserving the overall sum of synaptic displacements Σ |*w*_*i*_| as shown in equations 32 and 33. Across all results, the value of *κ* never reached below 1.0.

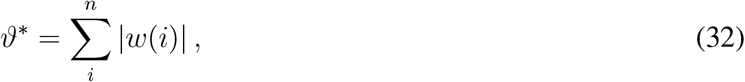

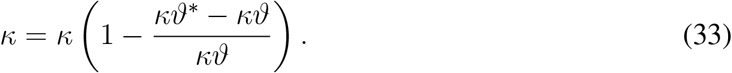

After update of the homeostatic parameter *κ*, we also update the sum of displacements *ϑ → ϑ*^*∗*^ .

### Simulations of reward-place association

Place information and reward-related information are represented by Poisson spike trains along a 200-cm-long treadmill. Specifically, place-informing Poisson units have place fields defined by Gaussian distributions, where their means are placed every 20 cm and identical variances are 10 cm. Note that the boundary place fields have means 0 and 200 cm. ‘Reward’ is delivered at a fixed location on the treadmill, but results are independent of the chosen reward location. Each trial is 180 seconds long with each lap lasting about 5 to 6 seconds. We deliver rewards between 20 to 100 seconds. Place-coding excitatory and inhibitory neurons and reward-coding VTA neurons show different minimum and maximum response frequencies. The values of these and other parameters are depicted in Table 3.

### Few-shot learning in networks of pre-existing assemblies

Our network results include preexisting assemblies with assembly sizes defined by a Gaussian distribution of mean = *μ* = 18 and standard deviation *σ* = 3. The cells in these networks had parameters as in Spike trace-based model from Table 1 with different capacitance for excitatory and inhibitory cells to increase the firing rate of inhibitory units. The rest of the parameters and their values are depicted in Table 3. In our analysis regarding weakly and strongly coupled assemblies(Fig. 7D), we picked the six strongest activated assemblies across the trial simulation for given salient patterns on each network. We also conditioned the response,in terms of spike count relative to assembly size, of these assemblies to be above 1. Any assembly that spiked at least a number of spikes equal to the assembly size at the last given pattern presentation is thus considered in the analysis. This favors the weakly coupled pre-existing assemblies as their activity is much lower than in the strongly coupled case. We consider that this is appropriate as small groups (2-3 cells) within weakly coupled assemblies may be responsive to a given salient pattern but the assembly itself has yet to develop a group response.

---

**Supplementary Figure 1:**
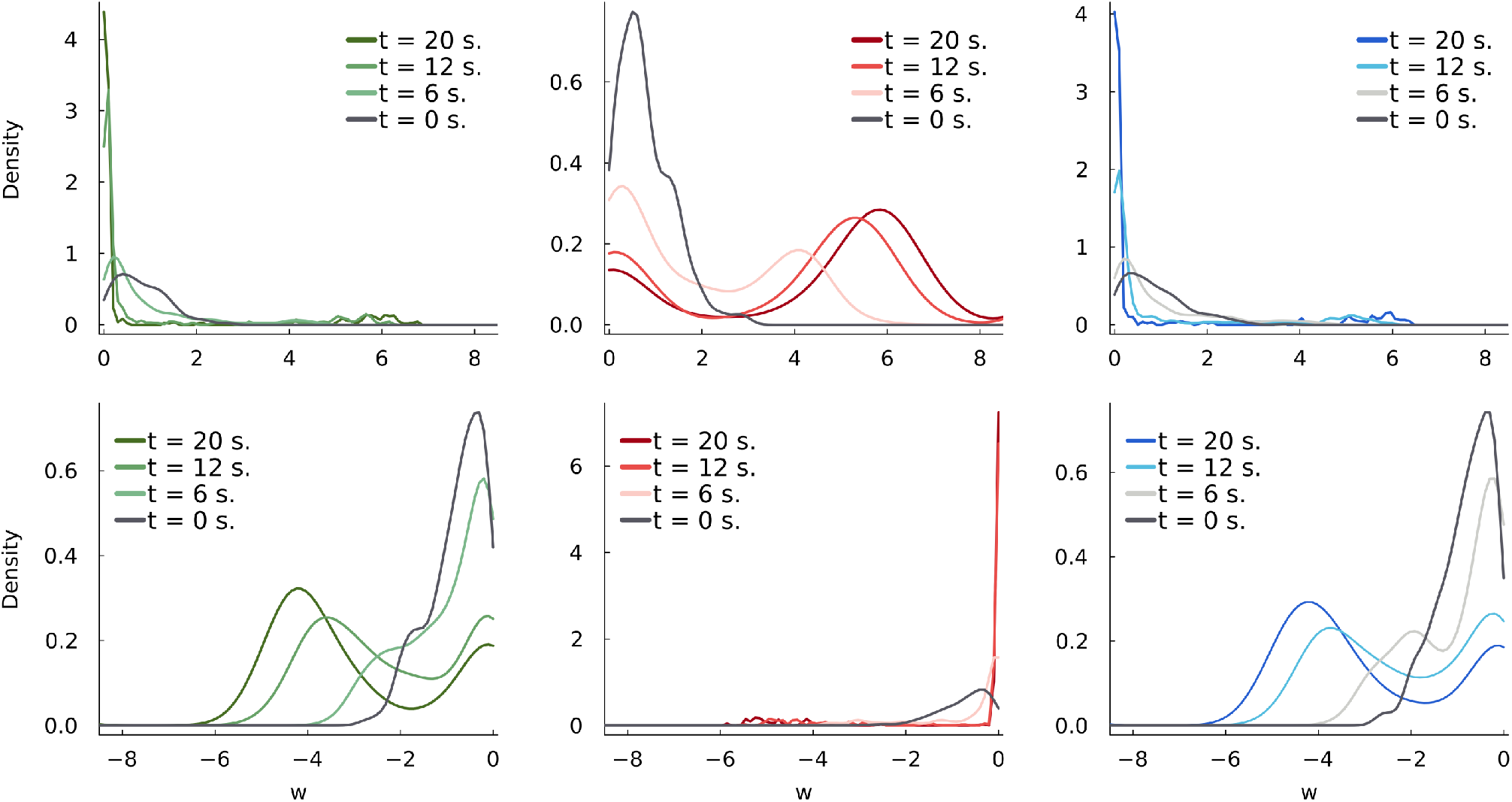
Kernel density estimation of excitatory and inhibitory synaptic weight *w* distributions for red tuned trial (both models yield same results). Colored lines from estimations at *t* = 0, 6, 12, 20 seconds of a single trial. (**top row**) Distributions for excitatory synapses pooled for red, green, and blue patterns. Across time, red pattern pooled excitatory synapses become bimodal with a large amount of strong synapses. (**bottom row**) Distributions for inhibitory synapses pooled for red, green, and blue patterns. Across time, blue and green pooled inhibitory synapses become bimodal exhibiting strong inhibition with blue and green pattern presentations relative to red pooled inhibitory synapses. Selectivity to a single pattern is hinted as both dense excitatory links for red pattern as well as dense inhibitory links for green and blue pooled synapses.

**Supplementary Figure 2:**
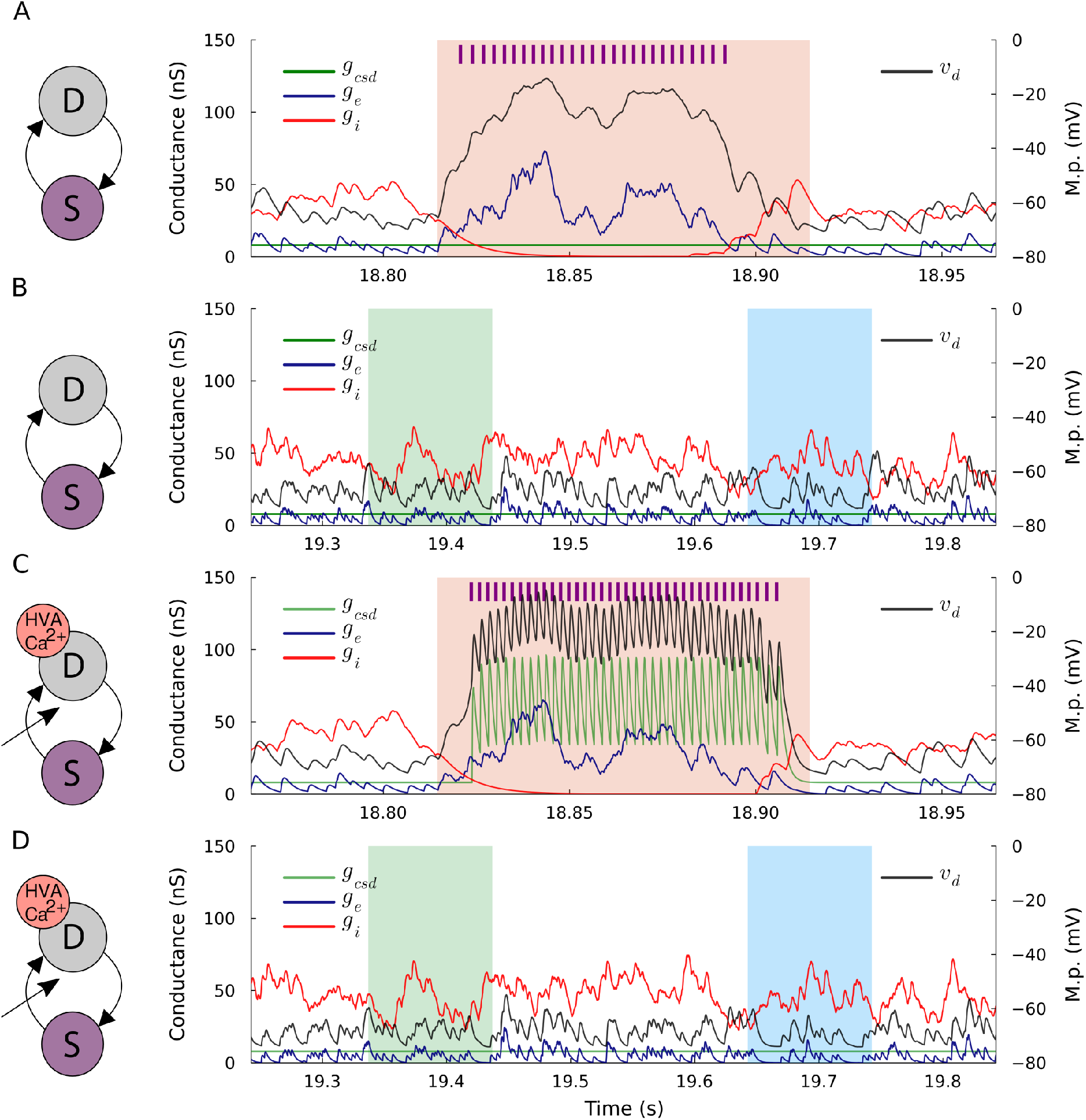
Example time course of conductance traces during pattern presentation for spike trace-based and calcium-based models. The corresponding model layout on the left of each time course plot. Red pattern tuned trial. (**A**) Spike trace-based model conductances (blue: excitatory, red: inhibitory, green: somato-dendritic) and dendritic membrane trace (black; axis on the right) during red pattern presentation at the end of the simulation. Vertical lines on top represent model output spikes. Inhibitory conductance plummets as an effect of learning while excitatory conductance remains high during red pattern presentation. (**B**) Same as above but for green and blue pattern presentations. Inhibitory conductance is high relative to excitatory conductance. (**C**) Same as in A but for calcium-based model conductances. *g*_*csd*_ is transiently boosted on each somatic spike to enhance dendritic membrane potential peaks. (**D**) Same as B but for the calcium-based model.

**Supplementary Figure 3:**
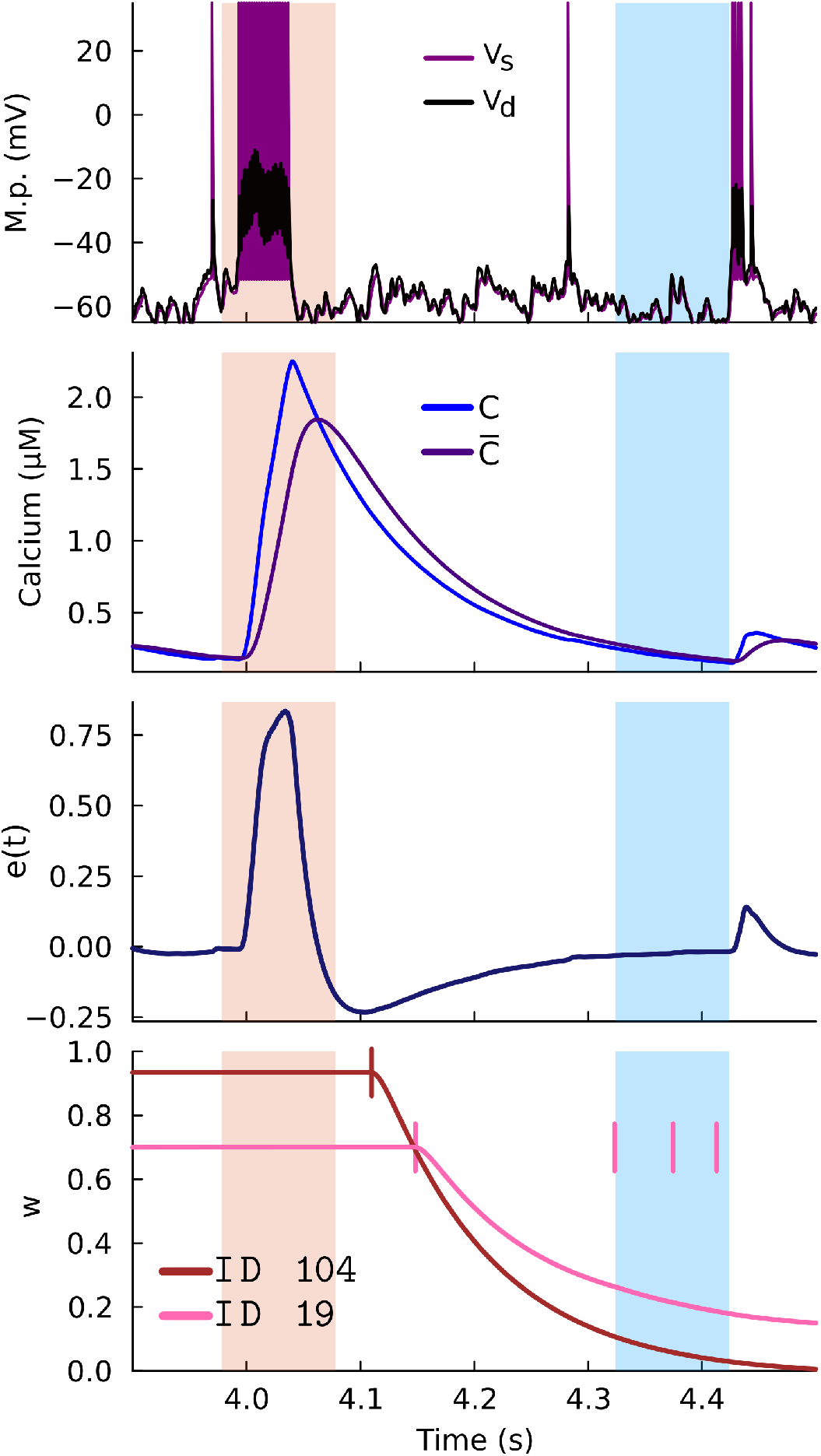
Example trial simulation for strong LTD induction to non-related inputs for calcium-based variant rule during pattern tuning. (**first row**) Membrane potential traces (left y-axis) of somatic (purple) and dendritic (gray) compartments. (**second row**) Calcium *C*(*t*) (blue) and low-pass filtered *C*(*t*) (midnight blue) traces. (**third row**) Plasticity induction error signal *e*(*t*). (**fourth row**) Synaptic weight changes Δ*w* of input IDs #109 (scarlet) and #19 (pink) traces across time. Vertical marker indicating spike time of input. Strong potentiation is followed by a long-lasting LTD phase yielding a negative phase of *e*(*t*) after the preferred pattern. Note that spikes of ID #19 (pink) occurring during the blue pattern do not affect plasticity induction of *w*_19_ as the sample of *e*(*t*) at given spike times is low (trend remains the same).

**Supplementary Figure 4:**
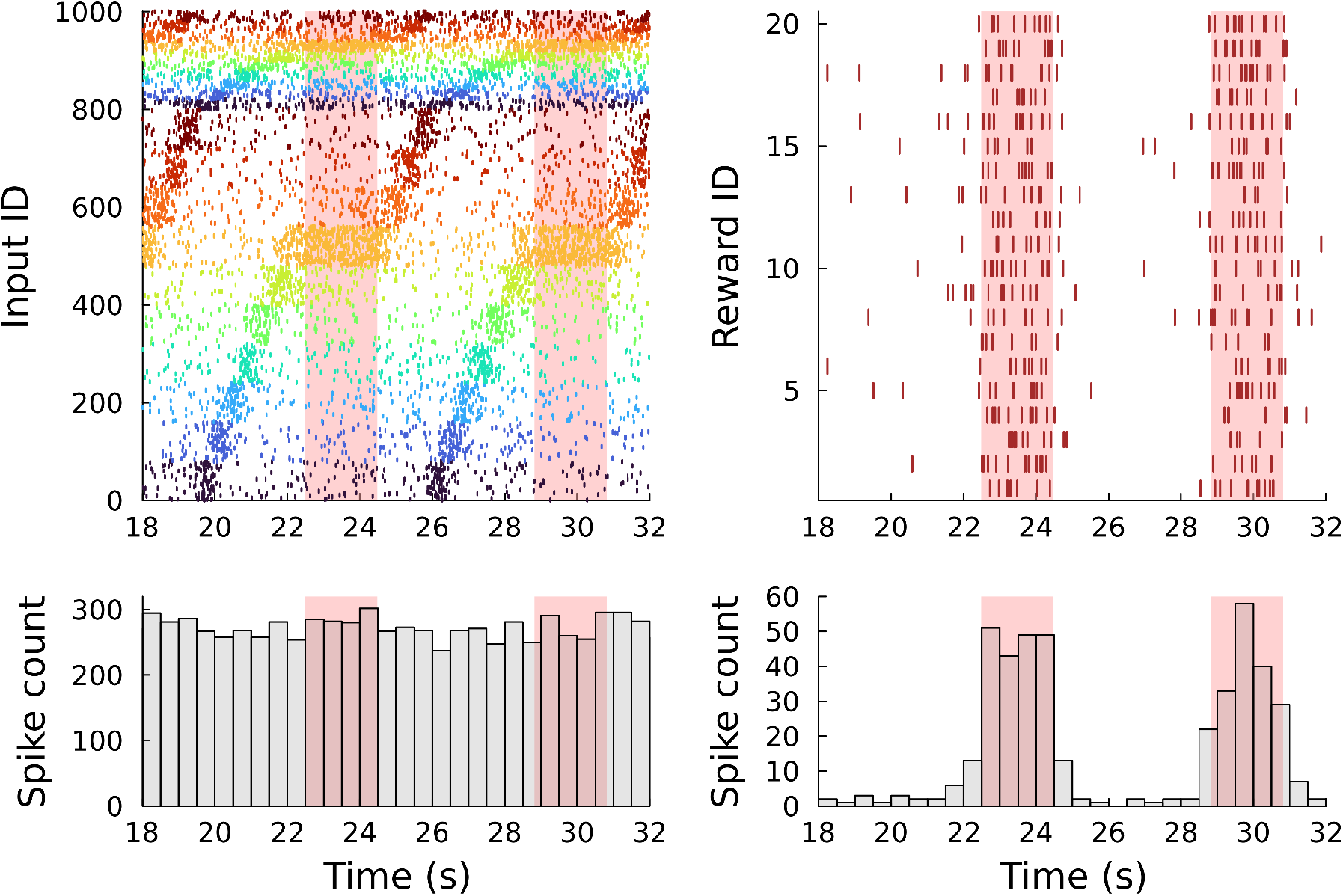
Raster plot and spike count plot of place field (PF) Poisson responses and reward cue inputs. (**left**) Raster plot of PF inputs (n=1000). Colored vertical bars are spike times as in Figure 6. Red colored spans represent `REWARD` states. Inhibitory inputs also have place field-like response curves. Spike count plot of all inputs binned every 500 ms. (**right**) Same as left but for reward cue inputs (n=20) during a single trial. Spiking frequency increases to *f*_*rew,max*_ during `REWARD` states.

**Supplementary Figure 5:**
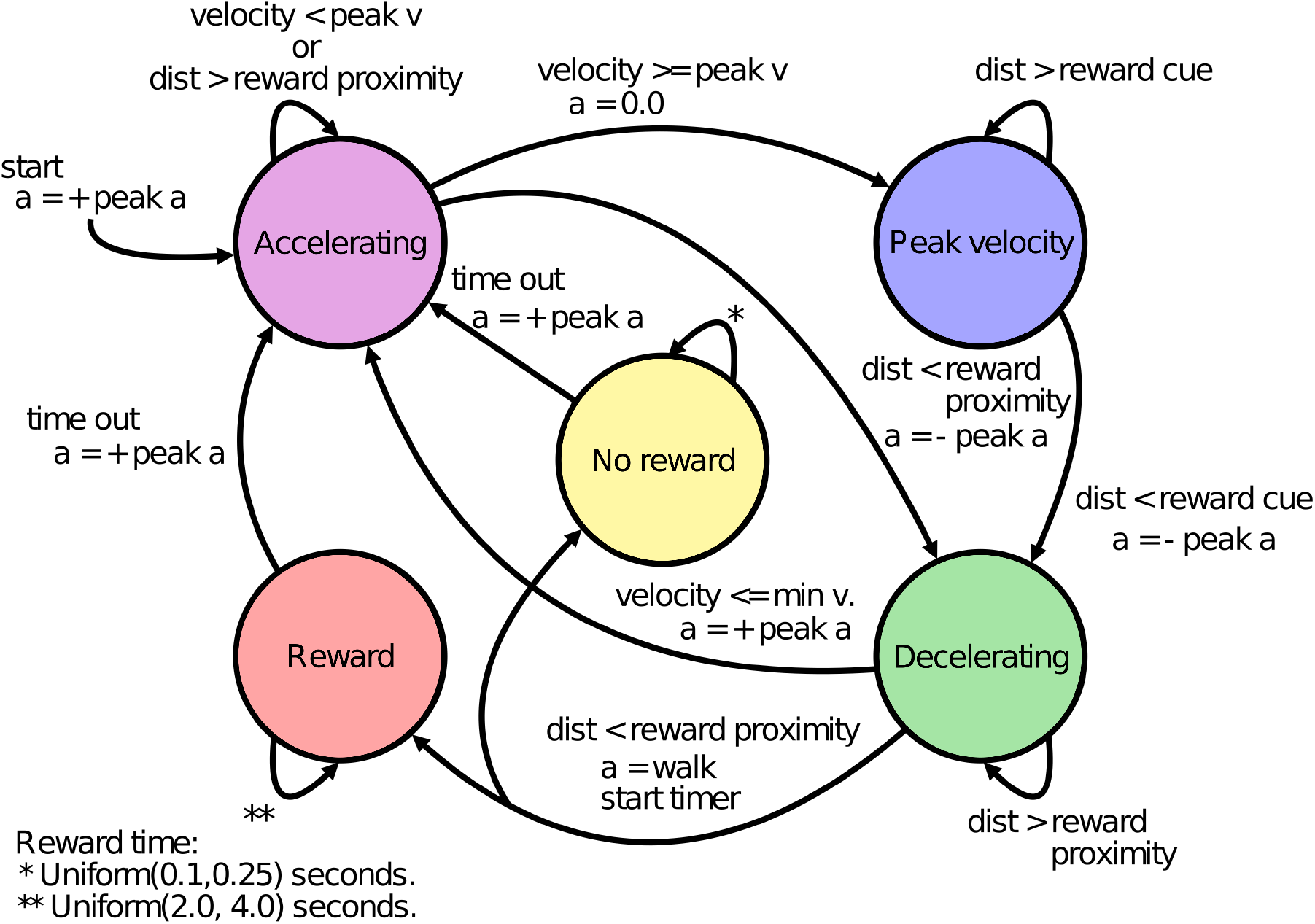
Behavior-mimicking finite state machine for reward-place association framework. FSM illustrated with transition conditions as `if` statements. a: acceleration, v: velocity, dist: absolute distance to reward. `NO REWARD` and `REWARD` states times are sampled from Uniform distributions. Random licking (`NO REWARD`) state considers proximity to reward location. Note that the colors of states are matched with the vertical color spans on the upper panel of Figure 6B

